# Modes of programmed macrophage cell death govern outcome of cutaneous wound healing

**DOI:** 10.64898/2026.03.19.712831

**Authors:** Louise Injarabian, Nils Reiche, Sebastian Willenborg, Daniela Welcker, Yunfan Bai, Eva Schönenberg, David E Sanin, Jovan Tanevski, Manolis Pasparakis, Hamid Kashkar, Sabine A Eming

## Abstract

Misregulation of tissue repair programs can severely compromise repair outcome. Timely clearance of inflammatory macrophages through regulated cell death is a prerequisite for resolution of inflammation and successful repair. How different modes of macrophage regulated cell death regulate repair and direct healing outcome remains unclear. Using inducible genetic models to trigger macrophage necroptosis (FADD^iMKO^) or enhance apoptosis of macrophages (cIAP1^iMKO^cIAP2^-/-^), we reveal opposing effects on the architecture of the wound tissue. Macrophage necroptosis profoundly disrupted tissue organization: FADD^iMKO^ wounds exhibited reduced numbers of reparative (IL-4Rα^⁺^Ly6C^low^) macrophages, delayed inflammatory resolution, accompanied by a hemorrhagic granulation tissue and reduced myofibroblast differentiation in the mid-phase of repair. In striking contrast, increased apoptosis preserved fundamental tissue architecture and vascular integrity, and reduced scar formation. Furthermore, single-cell transcriptomics demonstrated that macrophage necroptosis disrupts tissue-wide cellular communication networks essential for coordinated repair, in particular macrophage-fibroblast crosstalk. These findings establish that wound healing quality depends on both mode and rate of macrophage programmed death, providing a framework for therapeutic targeting of macrophages in wound healing disorders and fibrotic diseases.

## INTRODUCTION

The quality of tissue repair fundamentally determines clinical outcomes of wound healing, fibrotic diseases, and tissue regeneration. While acute wounds typically heal within weeks through coordinated cellular responses, repair is hampered by persistent inflammation and disorganized tissue architecture in chronic wounds (Eming *et al*, 2014). Similarly, maladaptive repair can lead to fibrosis of multiple organs including lung, liver, kidney, and skin, characterized by excessive extracellular matrix deposition and tissue dysfunction (Wynn & Vannella, 2016). Understanding the cellular mechanisms that determine whether damaged tissue regenerates its original functional architecture or progresses to chronic inflammation or fibrosis represents a critical challenge.

Macrophages are major orchestrators of tissue repair. Blood monocytes recruited to sites of injury differentiate into macrophages that exhibit remarkable functional plasticity, transitioning from pro-inflammatory Ly6C^high^ phenotypes during early inflammatory phases of repair toward pro-reparative Ly6Cl^ow^ phenotypes a later stage. The latter cells coordinate angiogenesis, extracellular matrix remodeling, and neutrophil clearance through efferocytosis (Lucas *et al*, 2010; Willenborg *et al*, 2012; Li *et al*, 2022). Beyond phenotypic switching, timely clearance of inflammatory macrophages from wound sites through regulated cell death (RCD) is essential for resolution of inflammation and restoration of tissue homeostasis (Injarabian *et al*, 2024; Pang *et al*, 2021). However, how macrophage cell death is temporally regulated during repair, and, critically, whether distinct modes of RCD differentially impact tissue architecture and healing quality, remains to be defined.

Emerging evidence reveals that different forms of RCD have context-dependent effects on tissue homeostasis and pathology (Blériot *et al*, 2015; Lee *et al*, 2023; Pasparakis & Vandenabeele, 2015). Necroptosis, a regulated form of inflammatory cell death characterized by plasma membrane rupture and release of damage-associated molecular patterns (DAMPs), contrasts sharply with apoptosis, traditionally viewed as immunologically “silent” (Galluzzi *et al*, 2018). Yet, in skeletal muscle injury, myocyte necroptosis promotes regeneration by stimulating stem cell proliferation (Zhou *et al*, 2020), whereas in liver, Kupffer cell necroptosis orchestrates type-2 immune responses (Blériot *et al*, 2015). Conversely, excessive apoptosis has been implicated in autoinflammatory diseases including inflammatory bowel disease and graft-versus-host disease (Prado-Acosta *et al*, 2023; Chiou *et al*, 2024). These tissue-specific outcomes underscore that cell death mode, beyond mere loss of the dying population, shapes inflammatory and repair responses.

Through transcriptional profiling of wound macrophages, we recently identified that early injury macrophages exhibit a TNF-mediated cell death signature with elevated *Tnfa*, *Fadd*, and *Ripk3* expression (Sanin *et al*, 2022; Injarabian *et al*, 2024). We demonstrated that combined inhibition of RIPK3-mediated necroptosis and FADD-mediated apoptosis in macrophages profoundly delays wound healing, with FADD deficiency primarily responsible for impaired repair (Injarabian *et al*, 2024). RIPK3 deficiency alone did not noticeably alter healing kinetics, raising critical questions: how does macrophage necroptosis impact tissue architecture and wound healing quality? Does apoptosis compromise tissue quality if in excess? These questions have direct therapeutic implications, as necroptosis inhibitors (targeting RIPK1/RIPK3) and apoptosis modulators (targeting BCL-2 family proteins) are under clinical investigation for inflammatory diseases.

To address these questions, we generated complementary genetic models to dissect effects of each death mode independently. We created mice with inducible, macrophage-specific deletion of *Fadd* to trigger necroptosis (Fadd^fl/fl^ Cx3cr1^CreER/wt^, FADD^iMKO^) as cells react to inactivation of the FADD-dependent apoptosis pathway by initiating necroptosis (Bonnet *et al*, 2011). For induced apoptosis of macrophages, we combined deletion of *Birc2* and *Birc3* to enhance apoptosis (Birc2^fl/fl^Birc3^-/-^Cx3cr1^CreER/wt^, cIAP1^iMKO^cIAP2^-/-^) (Wong *et al*, 2014). We hypothesized that inflammatory necroptosis would disrupt tissue organization, while apoptosis, being less inflammatory, would be better tolerated.

We indeed observed opposing effects of the two modes of RCD: macrophage necroptosis disrupted tissue architecture and wound healing quality characterized by a reduction of reparative macrophages (IL-4Rα^+^Ly6C^low^), hemorrhagic granulation tissue, reduced myofibroblast differentiation and delayed inflammation resolution. Single-cell transcriptomics demonstrated that macrophage necroptosis disrupts tissue-wide cellular communication networks essential for coordinated repair. In striking contrast, increased macrophage apoptosis reduced the number of wound macrophages, yet preserved fundamental tissue structure and vascular stability.

These results establish a framework for understanding how macrophage RCD mode determines repair outcomes, with direct implications for therapeutic targeting of cell death pathways in wound disorders and fibrotic diseases.

## RESULTS

### Genetic mouse models enabling induction of macrophage necroptosis or apoptosis during wound healing

To induce macrophage necroptosis or enhance apoptosis during wound healing without causing embryonic lethality or systemic inflammation, we intended to rely on tamoxifen-inducible *Cx3Cr1*^CreER^-mediated deletion of *Fadd* (to induce necroptosis) or *Birc2* (encoding cIAP1, on a *Birc3*^-/-^ background; to enhance apoptosis) (Yona *et al*, 2013; Wong *et al*, 2014).

To first obtain an estimate of the efficiency of Cre-mediated recombination we can achieve in the wound macrophage compartment of *Cx3Cr1^CreER^* mice, we crossed them to *R26^loxSTOPlox-^*^YFP^ Cre excision reporter mice as described in Yona et al. Upon Cre induction by intraperitoneal injection of tamoxifen daily for five consecutive days prior to wounding (**Fig. S1A**), we quantified YFP^+^ cells among CD45^+^CD11b^+^F4/80^+^ macrophages in wound tissue of *Cx3Cr1*^CreER^:*R26*^YFP^ (YFP-Cx3Cr1^CreER^) animals. Immunofluorescence analysis of frozen wound tissue samples revealed YFP^+^F4/80^+^ double-positive cells within granulation tissue at 2, 4 and 7 dpi (**Fig. 1A**). Flow cytometric analysis of single cell suspensions of wound tissue indicated substantial recombination efficiency in macrophages during the inflammatory phase with about 20% of total wound macrophages expressing the reporter at days 2 and 4 post injury (dpi). (**Fig. 1B-D**). At later stages, 7 days dpi, the fraction of reporter-expressing macrophages and their absolute numbers dropped, suggesting that initially labeled macrophages are lost from the wounds during the repair process. Analysis of kidney macrophages at 7 dpi in *Cx3Cr1*^CreER^;*R26*^YFP^ reporter mice showed high labeling efficiency throughout skin wound healing (**Fig. 1E, F**), consistent with published high recombination efficiency of kidney macrophages in this model (Yona *et al*, 2013).

**Figure 1.**
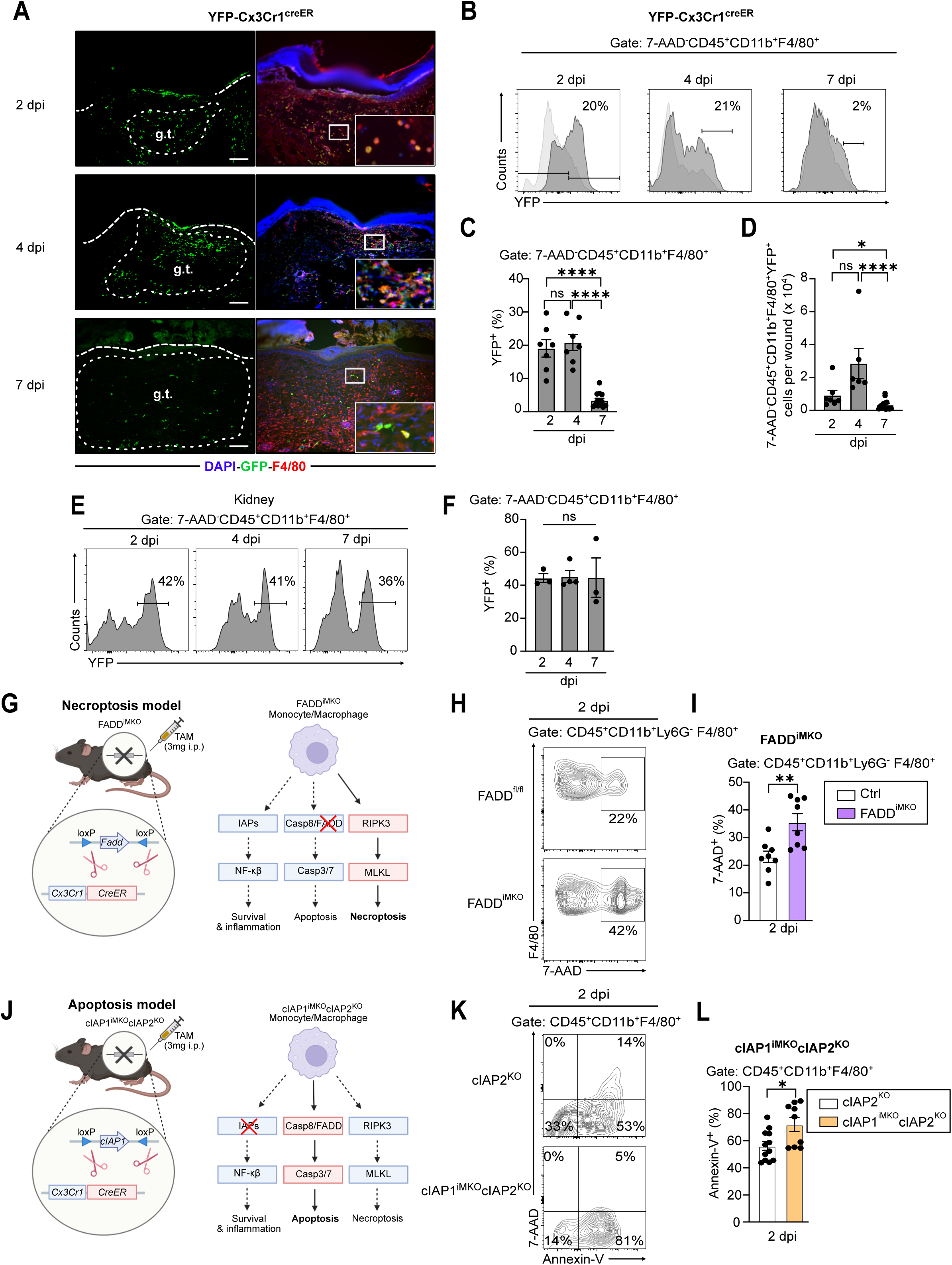
Inducible genetic models to trigger distinct modes of macrophage cell death during wound healing. **(A)** Representative immunofluorescence images of cryosections from YFP-Cx3Cr1^CreER^ reporter mice at 2, 4, and 7 days post-injury (dpi), co-stained with anti-F4/80 (red) and anti-GFP (green) antibodies. DAPI (blue) marks nuclei. Small dotted lines delineate the granulation tissue (g.t.) and larger lines the epithelium. Scale bar = 100 μm. **(B)** Representative flow cytometry histograms showing YFP^+^ macrophages (gated on 7-AAD^−^CD45^+^CD11b^+^F4/80^+^) in wounds at 2, 4, and 7 dpi. **(C)** Quantification of relative (%) and **(D)** absolute YFP^+^F4/80^+^ macrophage numbers per wound across the healing time course. **(E)** Representative flow cytometry histograms showing YFP^+^ macrophages in the kidney at 2, 4, and 7 dpi. **(F)** Quantification of relative YFP^+^ kidney macrophage numbers (%) across the healing time course. **(G)** Schematic illustration of the FADD^iMKO^ mouse model showing tamoxifen-inducible, macrophage-specific deletion of *Fadd* via Cx3cr1^CreER^-mediated recombination, and the resulting pathway shift from survival/apoptosis toward necroptosis (RIPK3–MLKL signaling). **(H)** Representative flow cytometry plots showing 7-AAD^+^ (dead) macrophages (gated on CD45^+^CD11b^+^Ly6G^−^F4/80^+^) in FADD^iMKO^ and control wounds at 2 dpi. **(I)** Quantification of 7-AAD^+^ macrophage frequency (%) in FADD^iMKO^ versus control wounds at 2 dpi. **(J)** Schematic illustration of the cIAP1^iMKO^cIAP2^KO^ mouse model showing tamoxifen-inducible deletion of *Birc2* (cIAP1) on a *Birc3*^−/−^ background, and the resulting pathway shift toward enhanced apoptosis (Casp3/7 activation). **(K)** Representative flow cytometry scatter plots showing Annexin-V and 7-AAD staining in macrophages (gated on CD45^+^CD11b^+^F4/80^+^) from cIAP2^KO^ control and cIAP1^iMKO^cIAP2^KO^ wounds at 2 dpi. Quadrants indicate viable (Annexin-V^−^7-AAD^−^), early apoptotic (Annexin-V^+^7-AAD^−^), late apoptotic (Annexin-V^+^7-AAD^+^), and secondary necrotic (Annexin-V^−^7-AAD^+^) cells. **(L)** Quantification of total apoptotic macrophage frequency (Annexin-V^+^7-AAD^−^ plus Annexin-V^+^7-AAD^+^; % of CD45^+^CD11b^+^F4/80^+^) in cIAP1^iMKO^cIAP2^-/-^ versus cIAP2^-/-^ control wounds at 2 dpi. Data are presented as mean ± SEM. Statistical significance was determined by two-tailed Student’s t-test (*p < 0.05; **p < 0.01; ****p < 0.0001; ns, not significant). Each data point represents an individual wound. All experiments were performed at least twice independently with a minimum of 5 mice per condition.

Next, we generated *Fadd*^fl/fl^;*Cx3cr1*^CreER/wt^ (FADD^iMKO^) mice to induce macrophage necroptosis (**Fig. 1G**). To evaluate the efficiency of cell death induction, we analyzed single cell suspensions of wound tissue by flow cytometry. Analysis revealed significantly increased macrophage cell death (7-AAD^+^) in FADD^iMKO^ wounds when compared to controls at 2 dpi (**Fig. 1H, I**). Increased fractions of dead macrophages were also observed at 4 and 7 dpi, whereas no difference was observed at 14 dpi (**S1B**).

To inducibly trigger macrophage apoptosis, we generated *Birc2*^fl/fl^*Birc3*^-/-^*Cx3cr1*^CreER/wt^ (cIAP1^iMKO^cIAP2^-/-^) mice (**Fig. 1J**). Deletion of both cIAP1 and cIAP2 is necessary due to functional redundancy. Flow cytometric analysis revealed significantly increased fractions of apoptotic macrophages (7-AAD^-^Annexin-V^+^ + 7-AAD^+^Annexin-V^+^) in cIAP1^iMKO^cIAP2^-/-^wounds at 2 dpi when compared to controls (**Fig. 1K, L**).

Importantly, neither model caused substantial macrophage depletion as we observed roughly 20% increase of cell death in both models. Moreover, analysis of cell death in Ly6C^low^ versus Ly6C^high^ macrophage subpopulations using flow cytometry revealed no selective elimination of a specific subset (**Fig. S1E-H**). These findings establish that both models successfully induce distinct death modes without causing excessive macrophage loss, allowing us to assess effects of death mode rather than macrophage depletion.

### Macrophage necroptosis reduces reparative Ly6C^low^ macrophages and significantly disrupts tissue architecture in the mid-phase of repair

To determine how macrophage necroptosis impacts tissue repair and architecture, we subjected FADD^iMKO^ mice to excision skin injury. Morphologically, wounds appeared larger in FADD^iMKO^ mice when compared to controls at 7 dpi (**Fig. 2A**), suggesting an impaired wound healing response. To analyzed immune cell dynamics across the healing time course we subjected single cell suspensions of wound tissue at subsequent phases of repair to flow cytometry analysis. Analysis revealed decreased viable macrophage numbers in FADD^iMKO^ wounds at 2, 4, and 7 dpi, with recovery by 14 dpi (**Fig. 2A, B**)

**Figure 2.**
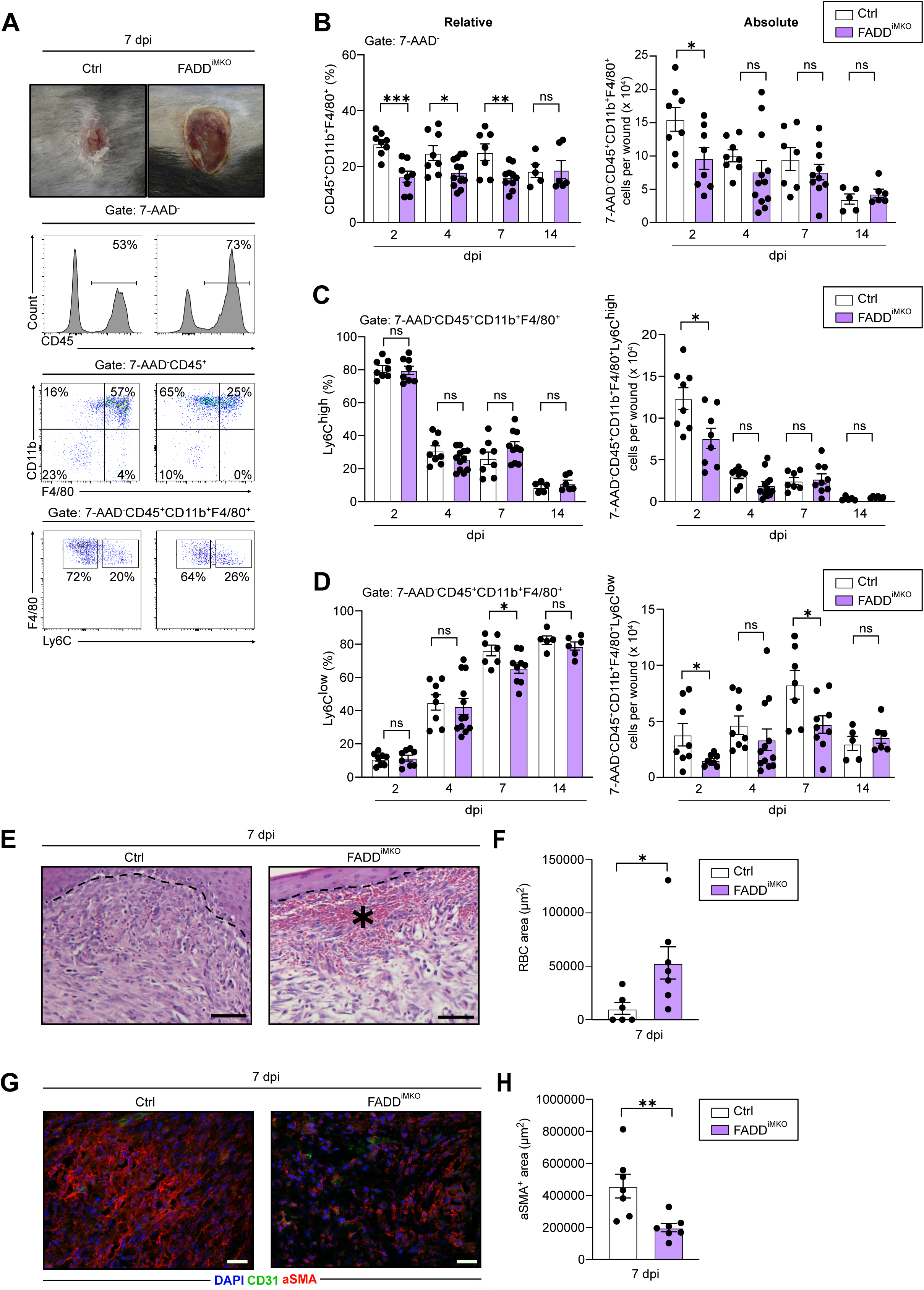
Macrophage necroptosis depletes Ly6C^low^ reparative macrophages and causes hemorrhagic tissue defects. **(A)** Representative macroscopic wound images (top) and flow cytometry histograms (bottom) showing viable macrophage gating (7-AAD^−^CD45^+^CD11b^+^F4/80^+^) and Ly6C^high^/Ly6C^low^ subpopulation identification in FADD^iMKO^ and control wounds at 7 dpi. **(B)** Quantification of relative (left) and absolute (right) viable macrophage numbers in FADD^iMKO^ versus control wounds across the healing time course (2, 4, 7, and 14 dpi). **(C)** Quantification of relative (left) and absolute (right) Ly6C^high^ pro-inflammatory macrophage numbers in FADD^iMKO^ versus control wounds across the healing time course. **(D)** Quantification of relative (left) and absolute (right) Ly6C^low^ reparative macrophage numbers in FADD^iMKO^ versus control wounds across the healing time course. **(E)** Representative hematoxylin and eosin (H&E)-stained histological sections of FADD^iMKO^ and control wounds at 7 dpi. Asterisk indicates hemorrhagic area. he, hyperproliferative epithelium; d, dermis; dWAT, dermal white adipose tissue; pc, panniculus carnosus; gt, granulation tissue. Scale bar = 50 μm. **(F)** Quantification of hemorrhagic area (red blood cell area, μm²) within the granulation tissue of FADD^iMKO^ versus control wounds at 7 dpi. **(G)** Representative immunofluorescence images of wound sections co-stained with anti-αSMA (red; myofibroblast marker), anti-CD31 (green; endothelial marker), and DAPI (blue) in FADD^iMKO^ and control mice at 7 dpi. Scale bar = 50 μm. **(H)** Quantification of αSMA^+^ area (μm²) within the granulation tissue of FADD^iMKO^ versus control wounds at 7 dpi. Data are presented as mean ± SEM. Statistical significance was determined by two-tailed Student’s t-test (*p < 0.05; **p < 0.01; ***p < 0.001; ns, not significant). Each data point represents an individual wound. All experiments were performed at least twice independently with a minimum of 5 mice per condition.

Analysis of macrophage subpopulations revealed that while Ly6C^high^ pro-inflammatory macrophages showed reduced absolute numbers at 2 dpi (**Fig. 2C**), Ly6C^low^ reparative macrophages exhibited profound depletion at both 2 dpi and 7 dpi (**Fig. 2D**). Because previous cell death analysis showed no increased cell death of Ly6C^low^ macrophages at 7 dpi (**Fig. S1G**), these findings suggest that macrophage necroptosis could impair the emergence of Ly6C^low^ macrophages through reduced reprogramming or proliferation.

Ly6C^low^ macrophages orchestrate granulation tissue maturation and vascularization (Rodero *et al*, 2013). To investigate the impact of reduced Ly6C^low^ macrophages on tissue architecture, we performed histomorphological analysis of FADD^iMKO^ and control wounds at 7 dpi and found no significant difference in total granulation tissue area (**Fig. S2A**). Strikingly, FADD^iMKO^ wounds exhibited extensive hemorrhages beneath the epithelium at 7 dpi (**Fig. 2E, F**). These hemorrhagic defects were accompanied by significantly reduced myofibroblast differentiation at 7 dpi (αSMA+ area: 200,000 ± 70,000 μm² vs 460,000 ± 20,000 μm²; p < 0.01) (**Fig. 2G, H**), indicating either delayed or impaired granulation tissue maturation. No significant differences were observed in overall vessel density, as CD31^+^ area within the granulation tissue remained comparable between the two conditions (**Fig. S2B**).

The hemorrhagic phenotype in FADD^iMKO^ mice phenocopied that observed upon macrophage depletion (Lucas *et al*, 2010) or myeloid-specific *Il4ra* deletion (Knipper *et al*, 2015). Because IL-4Rα is predominantly expressed by Ly6C^low^ macrophages in wounds (data from Sanin et al. 2022), we hypothesized that a reduction in Ly6C^low^ macrophages could impair IL-4Rα-dependent vascular stabilization. Immunofluorescence analysis confirmed strongly reduced IL-4Rα^+^ macrophages in FADD^iMKO^ compared to control wounds at 7 dpi (**Fig. S2C, D**). These findings suggest that macrophage necroptosis could compromise vascular integrity, likely through the reduction of IL-4Rα-dependent vessel-protective signals derived from Ly6C^low^ macrophages.

### Macrophage necroptosis causes persistent neutrophil accumulation and impairs tissue-wide efferocytic capacity

Ly6C^low^ reparative macrophages are the principal orchestrators of neutrophil clearance through efferocytosis, a process essential for inflammatory resolution (Lucas *et al*, 2010; Bosurgi *et al*, 2017). Given the profound depletion of Ly6C^low^ macrophages and the hemorrhagic vascular instability observed in FADD^iMKO^ wounds (Fig. 2), we asked whether neutrophil clearance was compromised. Flow cytometry revealed comparable neutrophil numbers at 2 and 4 dpi between FADD^iMKO^ and control wounds. However, while control wounds cleared most neutrophils by day 7, this did not occur in FADD^iMKO^ wounds which maintained high neutrophil numbers. Neutrophils were largly absent from control and FADD^iMKO^ wounds by 14 dpi (**Fig. 3A-B**). Histological analysis confirmed elevated Ly6G^+^ neutrophil numbers within the granulation tissue at 7 dpi (**Fig. S3A, B**). The temporal pattern of normal early influx followed by mid-phase persistence and eventual resolution suggests impaired efferocytic clearance rather than enhanced recruitment through the leaky vasculature.

**Figure 3.**
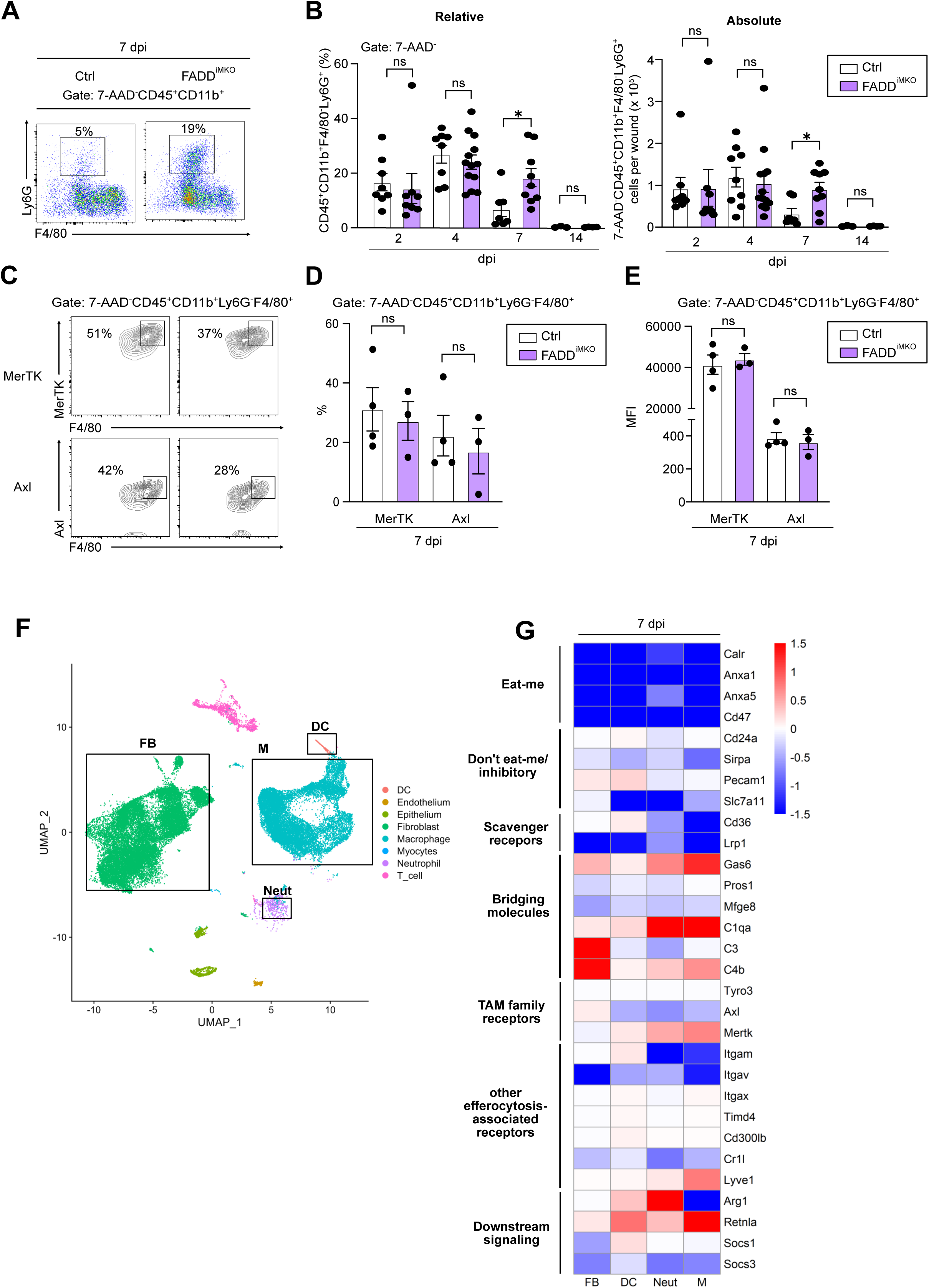
Macrophage necroptosis leads to persistent neutrophil infiltration and disrupts tissue-wide efferocytosis networks in FADD^iMKO^ wounds. **(A)** Representative flow cytometry scatter plots showing neutrophils (gated on 7-AAD^−^CD45^+^CD11b^+^F4/80^−^Ly6G^+^) in FADD^iMKO^ and control wounds at 7 dpi. **(B)** Quantification of relative (left) and absolute (right) neutrophil numbers in FADD^iMKO^ versus control wounds across the healing time course (2, 4, 7, and 14 dpi). **(C)** Representative flow cytometry contour plots showing MerTK (top) and Axl (bottom) expression on wound macrophages (gated on 7-AAD^−^CD45^+^CD11b^+^Ly6G^−^F4/80^+^) in FADD^iMKO^ and control mice at 7 dpi. **(D)** Quantification of MerTK^+^ and Axl^+^ macrophage frequencies (%) in FADD^iMKO^ versus control wounds at 7 dpi. **(E)** Quantification of MerTK and Axl mean fluorescence intensity (MFI) in wound macrophages from FADD^iMKO^ versus control mice at 7 dpi. Each data point represents an individual wound. All experiments were performed with at least 3 mice per condition. **(F)** Heatmap showing log2 fold-change (FADD^iMKO^ vs. control) of efferocytosis-related genes grouped by functional category (“eat-me” signals, “don’t-eat-me”/inhibitory signals, scavenger receptors, bridging molecules, TAM family receptors, other efferocytosis-associated receptors, and downstream signaling) across fibroblasts (FB), dendritic cells (DC), neutrophils (N), and macrophages (M) at 7 dpi from scRNAseq data. Color scale indicates log2FC (red, upregulated; blue, downregulated). Data are presented as mean ± SEM. Statistical significance was determined by two-tailed Student’s t-test (*p < 0.05; ns, not significant).

To determine whether macrophage-intrinsic efferocytic receptor expression was altered, we assessed surface levels of MerTK and Axl on wound macrophages at 7 dpi by flow cytometry. No significant differences were detected between FADD^iMKO^ and control macrophages (**Fig. 3C, D**), indicating that per-cell receptor expression is maintained independent of necroptotic context.

To evaluate efferocytosis-related gene expression across all wound cell types, we performed single-cell RNA sequencing (scRNAseq) (see **Fig. 4, Fig. S4A, B** and Methods for dataset description). We analyzed differential expression of genes governing apoptotic cell recognition, including eat-me signals, don’t-eat-me signals, recognition receptors, and engulfment machinery, across fibroblasts, dendritic cells, neutrophils, and macrophages at 7 dpi (**Fig. 3F**), following framework described by Justynski et al. (Justynski *et al*, 2023). Analysis revealed a broad, cell-type-spanning suppression of eat-me signals in FADD^iMKO^ wounds. Paradoxically, the don’t-eat-me signal *Cd47* was also downregulated across cell types, suggesting a global dampening of apoptotic cell-sensing programs rather than a selective shift in the phagocytic balance. Within macrophages, recognition receptors showed divergent transcriptional changes: *Axl* was significantly downregulated while *MerTK* was upregulated, a divergence that likely reflects post-transcriptional stabilization of MerTK protein consistent with the unchanged surface expression detected by flow cytometry. The scavenger receptor *Lrp1* and the integrin *Itgav*, both involved in apoptotic cell tethering and recognition, were reduced. Together, these transcriptional changes suggest a reduced total efferocytic capacity across wound cells, providing a mechanistic basis for the persistent neutrophil accumulation observed at 7 dpi in FADD^iMKO^ wounds.

**Figure 4.**
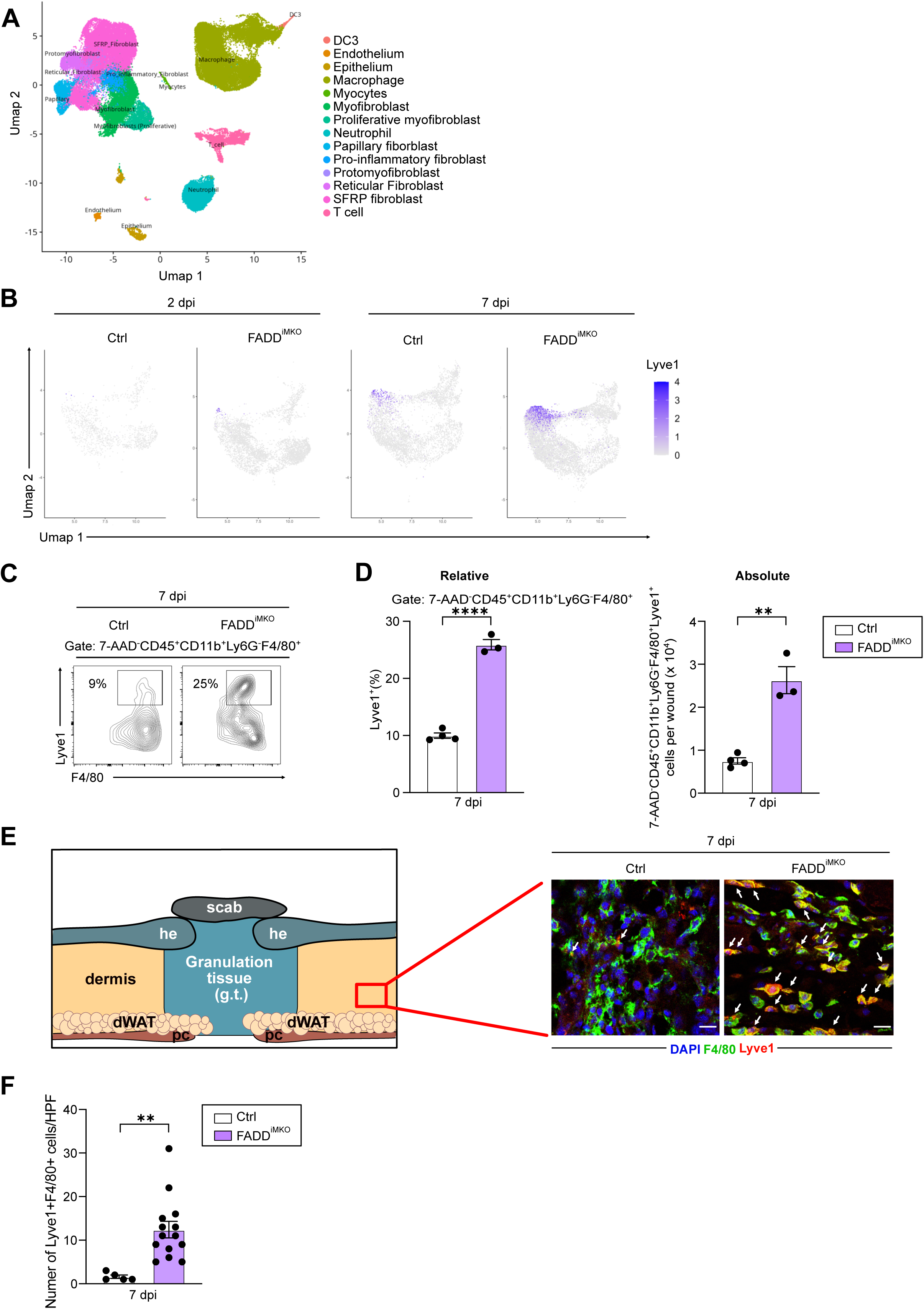
Macrophage necroptosis drives expansion of Lyve1^+^ macrophages in unwounded skin adjacent to wounds. **(A)** UMAP visualization of the integrated scRNAseq dataset from FADD^iMKO^ and control wounds, with cell types annotated by color. Cell populations include: DC3, Endothelium, Epithelium, Macrophage, Myocytes, Myofibroblast, Proliferative myofibroblast, Neutrophil, Papillary fibroblast, Pro-inflammatory fibroblast, Protomyofibroblast, Reticular fibroblast, SFRP fibroblast, and T cell. **(B)** Feature plots showing *Lyve1* expression (color scale: low = grey, high = dark blue) in the macrophage cluster from FADD^iMKO^ and control wounds at 2 and 7 dpi. **(C)** Representative flow cytometry contour plots showing Lyve1^+^ macrophages (gated on 7-AAD^−^CD45^+^CD11b^+^Ly6G^−^F4/80^+^) in FADD^iMKO^ and control wounds at 7 dpi. **(D)** Quantification of relative (left) and absolute (right) Lyve1^+^ macrophage numbers in FADD^iMKO^ versus control wounds at 7 dpi. **(E)** Schematic diagram of wound tissue anatomy (left) indicating the imaging region (red box) analyzed by immunofluorescence. Representative immunofluorescence images (right) of wound sections from control and FADD^iMKO^ mice at 7 dpi co-stained with anti-F4/80 (green), anti-Lyve1 (red), and DAPI (blue). Arrows indicate F4/80^+^Lyve1^+^ double-positive macrophages in the dermis adjacent to the wound. he, hyperproliferative epithelium; g.t., granulation tissue; dWAT, dermal white adipose tissue; pc, panniculus carnosus. Scale bar = 20 μm. **(F)** Quantification of F4/80^+^Lyve1^+^ macrophages per high-power field (HPF) in unwounded skin adjacent to wounds in FADD^iMKO^ versus control mice at 7 dpi. Data are presented as mean ± SEM. Statistical significance was determined by two-tailed Student’s t-test (**p < 0.01; ****p < 0.0001). Each dot represents an individual wound. All experiments were performed at least twice independently with a minimum of 5 mice per condition.

Interestingly, we observed an unexpected transcriptional signature in FADD^iMKO^ wound macrophages: elevated expression of *Lyve1* and *Retnla*, markers associated with vessel-associated, tissue-resident macrophages with high efferocytic and vascular-stabilizing capacity. This observation raised the question of whether a Lyve1^+^ macrophage subpopulation expands in response to necroptosis-driven vascular damage and whether such expansion is sufficient to rescue the tissue defects observed in FADD^iMKO^ wounds.

### Macrophage necroptosis reshapes the wound transcriptional landscape and drives compensatory Lyve1⁺ macrophage expansion in unwounded skin

To characterize the transcriptional alterations induced by macrophage necroptosis at single-cell resolution, we performed scRNAseq on wound tissue from FADD^iMKO^ and control mice at 2 and 7 dpi. Additionally, we used unwounded skin as a baseline control. After quality filtering and doublet removal (see Methods), we identified 14 transcriptionally distinct cell populations, annotated as dendritic cells (DC3), endothelium, epithelium, macrophages, neutrophils, myocytes, myofibroblasts, proliferative myofibroblast, papillary fibroblasts, pro-inflammatory fibroblasts, protomyofibroblasts, reticular fibroblasts, SFRP fibroblasts and T cells (**Fig. 4A; Fig. S5A-C**). Cell-type composition was broadly preserved between genotypes at 2 dpi; however, at 7 dpi, analysis revealed reduction in myofibroblasts (**Fig. S6**) and an increase in Lyve1 macrophages in FADD^iMKO^ wounds (**Fig.4B; Fig. S7A**), consistent with previous findings.

The prominent expression of *Lyve1* prompted us to investigate whether this reflected emergence of a distinct Lyve1^+^ macrophage subpopulation. Lyve1^+^ macrophages are vessel-associated tissue-resident macrophages known to stabilize microvasculature, promote angiogenesis, and support efferocytosis (Chakarov *et al*, 2019). Focused analysis of the macrophage cluster revealed that Lyve1^high^ cells were essentially absent at 2 dpi in both genotypes but robustly expanded in FADD^iMKO^ wounds as a distinct cluster at 7 dpi (**Fig. 4B; Fig. S7A**). Flow cytometry validated this finding (**Fig. 4C, D**).

Differential gene expression within the Lyve1^+^ macrophage cluster comparing FADD^iMKO^ to control wounds revealed 248 differentially expressed genes (adjusted p < 0.05; **Fig. S7B**). Upregulated genes included *Egfr* (log2FC =1.96, adjusted p < 0.01), consistent with macrophage proliferative responses, while the efferocytosis receptor *Cd36* was downregulated (log2FC = −1.86, adjusted p < 0.001) (**Fig. S7C**). Pathway analysis of genes upregulated in FADD^iMKO^ Lyve1^+^ macrophages revealed enrichment of antiviral and interferon-responsive programs (**Fig. S7D**), suggesting activation downstream of necroptotic danger signals, consistent with nucleic acids and HMGB1 released by necroptotic cells engaging pattern recognition receptors such as TLR3 and cGAS–STING.

To determine spatial distribution of Lyve1^+^ macrophages, we performed immunofluorescence analysis of wound tissue sections. Despite their numerical expansion, F4/80^+^Lyve1^+^ cells were predominantly localized to the unwounded skin adjacent to wounds rather than within the granulation tissue itself (12.4 ± 7 vs 1.6 ± 0.9 cells/HPF in adjacent skin, p < 0.01; **Fig. 4E, F**). To determine whether this spatial mismatch reflected injury-induced recruitment failure or pre-existing expansion of resident populations, we analyzed unwounded skin from FADD^iMKO^ and control mice. scRNAseq analysis confirmed increased Lyve1^+^ macrophages in FADD^iMKO^ skin prior to any wound stimulus (**Fig. S7E**), indicating that their expansion occurs before injury and most likely reflects a response to necroptosis in dermal-resident macrophages.

### Macrophage necroptosis replaces a reparative stromal-instructive signaling program with a damage-clearance communication state

The cellular and transcriptional phenotypes of FADD^iMKO^ wounds collectively suggested a breakdown in multicellular coordination. To test this hypothesis, we applied ligand–receptor interaction analysis (LIANA+;Dimitrov et al. 2024) to our annotated scRNAseq data to map the intercellular communication landscape of FADD^iMKO^ versus control wounds at 7 dpi.

Comparison of unique cell-cell interactions in controls and FADD^iMKO^ wounds revealed a fundamental remodeling of the signaling program. In control wounds, reticular fibroblast signaling was more frequent than in FADD^iMKO^ wounds (**Fig. 5A, B**). Furthermore, specific interaction analysis (ligand to receptor pairs) revealed that macrophages in control wounds engaged distinct fibroblast subpopulations through three integrated signaling modules (**Fig. 5C**). First, a lipid metabolic and scavenger receptor module: Apoe–Abca1/Lrp4 and Apoc2–Lrp1 signaling coordinated lipid handling between macrophages and papillary and reticular fibroblast subsets, alongside Thbs1–, S100a8–, and Saa1–Cd36 interactions spanning pro-inflammatory and reticular fibroblast populations. Second, a chemokine-mediated activation module: Ccl2–Ccr2, Ccl2–Ccr5, Ccl11–Ccr5, and Il16–Ccr5 axes directed inflammatory signals toward papillary, pro-inflammatory, and protomyofibroblast fibroblast states, consistent with macrophage-directed fibroblast priming. Third, a growth factor and morphogen module: Wnt5a–Ror1, Gas6–Tyro3, and multiple Fgf ligands (Fgf1, Fgf2, Fgf7, Fgf18) engaged Fgfr4-expressing fibroblast populations, supporting fibroblast proliferation, differentiation, and matrix remodeling. Together, control wound macrophages establish a multivalent stromal-instructive niche that coordinates fibroblast activation across spatially distinct dermal compartments.

**Figure 5.**
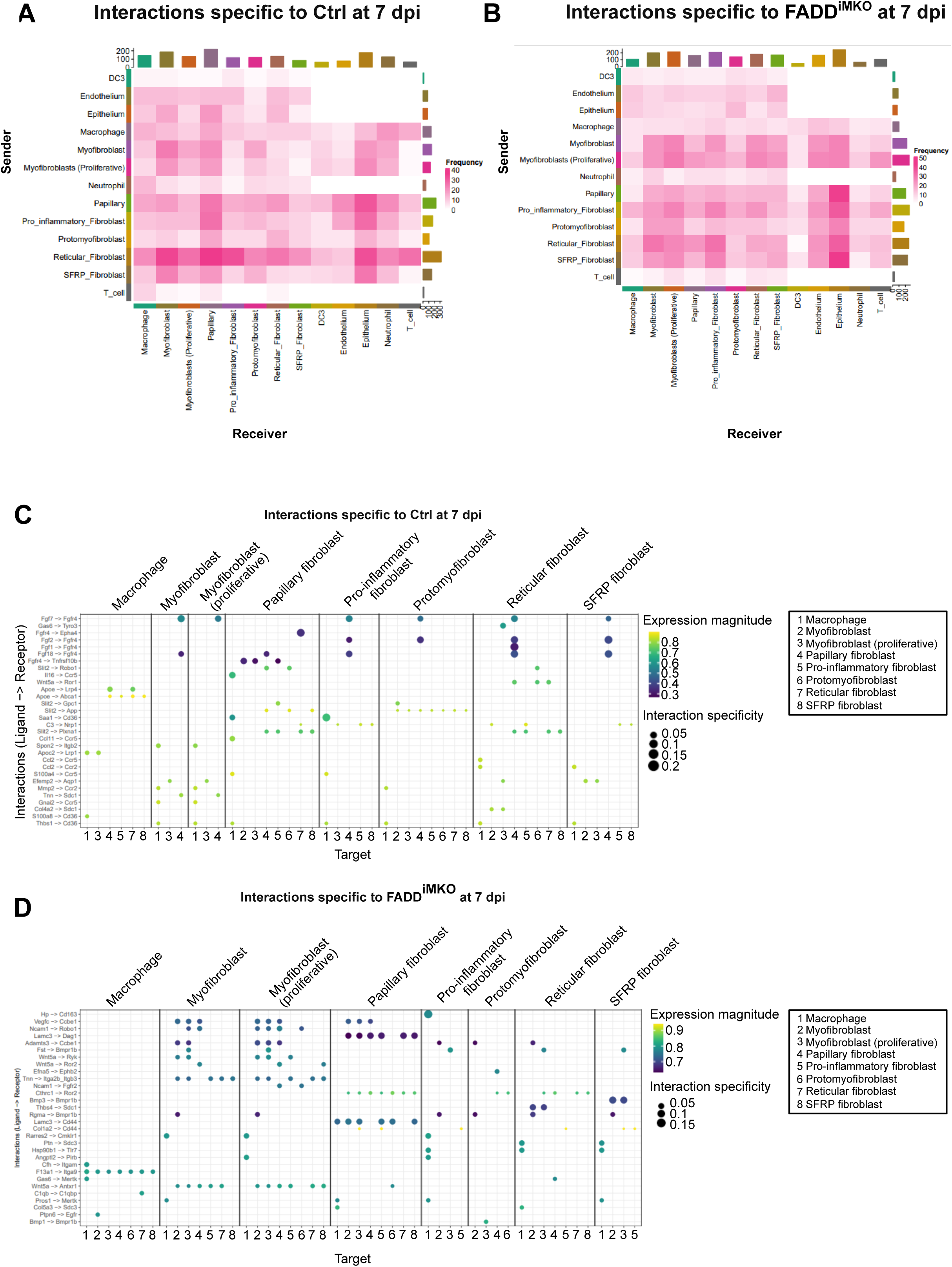
Macrophage necroptosis remodels tissue-wide cellular communication networks in FADD^iMKO^ wounds. **(A)** Heatmap showing the frequency of cell-type-specific ligand–receptor interactions unique in control wounds at 7 dpi, as inferred by LIANA+ analysis. Color intensity and cell-type color bars indicate interaction frequency. Rows, sending cell types; columns, receiving cell types. **(B)** Heatmap showing the frequency of cell-type-specific ligand–receptor interactions unique in FADD^iMKO^ wounds at 7 dpi. **(C)** Dot plot showing unique ligand–receptor interactions specific to control wounds at 7 dpi, with macrophages as sender cells and indicated fibroblast subpopulations as receivers. Dot color indicates expression magnitude (yellow–green scale); dot size indicates interaction specificity. Numbered target cell types are indicated in the legend. **(D)** Dot plot showing unique ligand–receptor interactions specific to FADD^iMKO^ wounds at 7 dpi, with macrophages as sender cells and indicated fibroblast subpopulations as receivers. Dot color indicates expression magnitude; dot size indicates interaction specificity. Numbered target cell types are indicated in the legend.

In FADD^iMKO^ wounds, this reparative program was replaced by a damage-clearance communication state (**Fig. 5D**). The most prominent shift was the emergence of hemoglobin-scavenging interactions, most notably Hp–Cd163, indicating macrophage engagement with extravasated erythrocyte-derived components, a signature of vascular instability and tissue hemorrhage. Concurrently, efferocytic and complement-associated interactions were enriched, including Gas6–MerTK, Pros1–MerTK, C1qb–C1qbp, and Cfh–Itgam, consistent with a macrophage program oriented toward debris clearance. Extracellular matrix interactions (Col1a2–Cd44, Lamc3–Cd44/Dag1, Thbs4–Sdc1) and aberrant morphogen signals through Bmp1/3–Bmpr1b and Wnt5a/Cthrc1–Ror2/Ryk further indicated disorganized matrix remodeling rather than coordinated stromal reconstitution. Thus, macrophage–fibroblast communication in FADD^iMKO^ wounds is defined by hemoglobin scavenging, complement activation, and disorganized ECM remodeling, a shift that is likely to compromise the fibroblast activation and myofibroblast differentiation programs required for normal granulation tissue maturation.

Notably, analogous communication network alterations were already apparent in unwounded FADD^iMKO^ skin (**Fig. S8A, B**). In control dermis, macrophages maintained homeostatic immune surveillance through MHC-II-mediated antigen-presentation interactions (H2-Aa/Ab1/Eb1/Dma–Cd4/Lag3), homeostatic chemokine signaling (Cxcl12–Cxcr4/Ackr3/Sdc4), and Csf2-dependent macrophage maintenance (Csf2–Csf1r/Csf2ra–Csf2rb), alongside fibroblast-derived matrix-sensing signals (Col1a1/Col1a2–Cd44, Has2–Cd44). By contrast, FADD^iMKO^ unwounded skin displayed chemokine-driven leukocyte recruitment axes (Ccl12–Ccr1/Ccr2/Ccr5), innate immune sensing pathways (C3–C3ar1, Cd14–Tlr1, Csf1–Sirpa), and enhanced integrin- and matrix-remodeling signals (Thy1–Itgam/Itgb2, Spon2–Itga4, Tnc–Itga2/Ptprz1, Wnt16–Fzd/Lrp6). This pre-existing shift from matrix-centered homeostatic surveillance toward immune-recruitment and stromal-remodeling indicates that macrophage necroptosis alters stromal–immune crosstalk basally and may lower the threshold for inflammatory amplification upon wounding.

Taken together, our analysis demonstrates that macrophage necroptosis does not simply reduce reparative macrophage numbers: it fundamentally remodels the intercellular signaling landscape, replacing a coordinated, growth factor-driven stromal-instructive program with a damage-response communication state that persists from homeostatic skin into the active wound repair phase

### Caspase-8 deletion in macrophages recapitulates FADD deletion defects during wound healing, confirming necroptosis as the primary driver

FADD serves dual roles in regulated cell death: as an adaptor for caspase-8-mediated apoptosis and as a negative regulator of RIPK3–MLKL-dependent necroptosis. To confirm that the wound healing defects observed in FADD^iMKO^ mice are attributable specifically to macrophage necroptosis rather than loss of FADD’s RCD-independent scaffolding functions, we generated an orthogonal necroptosis model through inducible, macrophage-specific deletion of caspase-8 (*Casp8*^fl/fl^*Cx3cr1*^CreER/wt^; Casp8^iMKO^). Caspase-8 deletion removes the suppression of RIPK3 activity, thereby directly inducing necroptosis in a manner molecularly distinct from FADD deletion.

Casp8^iMKO^ wounds recapitulated all cardinal defects observed in FADD^iMKO^ mice. Macrophage cell death was significantly increased at 2 dpi (**Fig. 6A, B**), supporting efficient induction of necroptosis. Neutrophil numbers were elevated at 7 dpi (**Fig. 6C, D**), and extensive hemorrhagic regions were present within the granulation tissue (**Fig. 6E, F**). The phenotypic concordance between two genetically distinct necroptosis models targeting different molecular checkpoints upstream of RIPK3 activation provides strong evidence that these tissue defects are driven by necroptosis specifically rather than by pleiotropic effects of FADD deletion.

**Figure 6.**
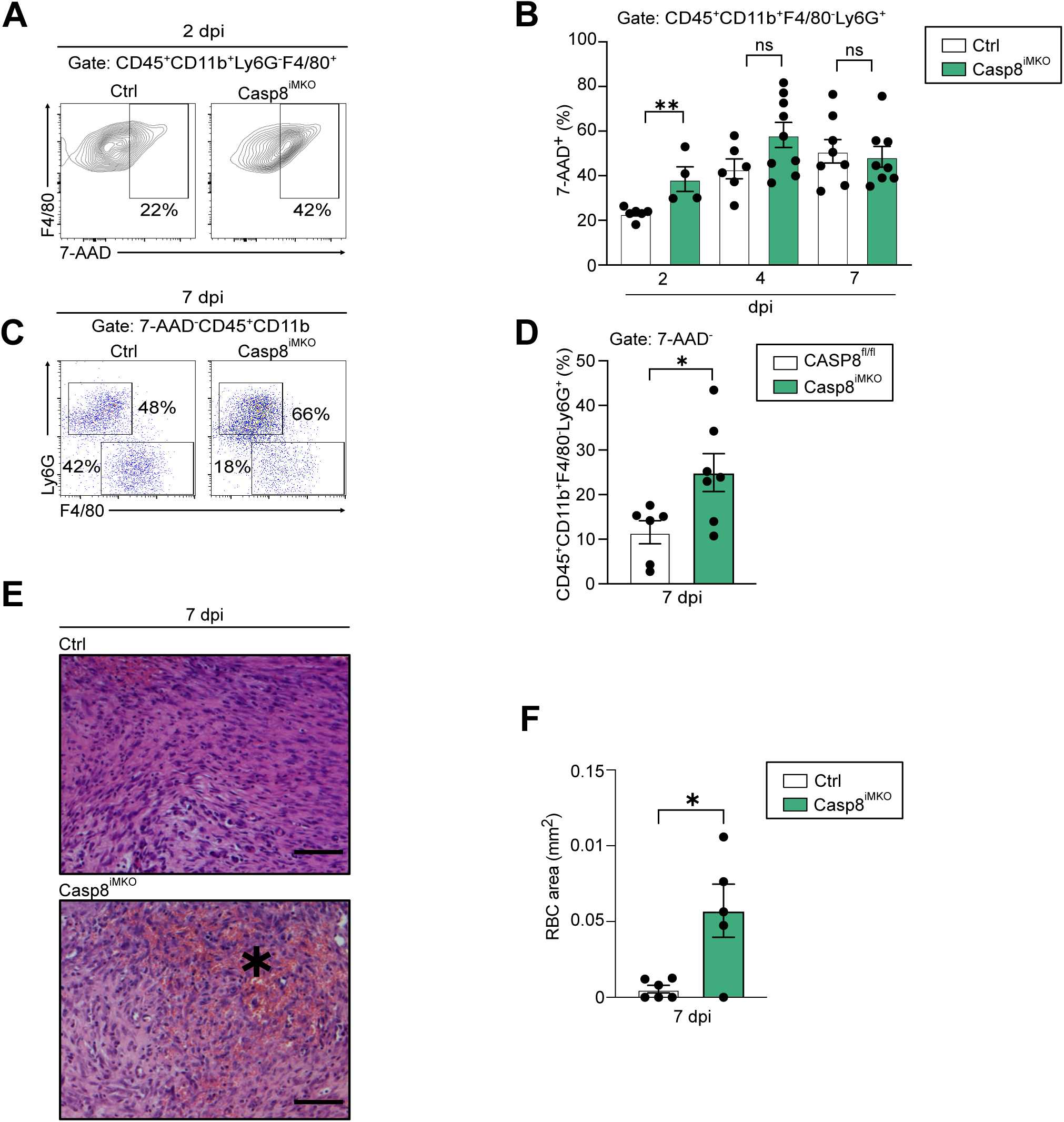
Caspase-8 deletion in macrophages phenocopies FADD^iMKO^ wound healing defects. **(A)** Representative flow cytometry scatter plots showing macrophage cell death (7-AAD^+^) gated on CD45^+^CD11b^+^Ly6G^−^F4/80^+^ cells in CASP8^iMKO^ and control wounds at 2 dpi. **(B)** Quantification of 7-AAD^+^ macrophage frequency (%) in CASP8^iMKO^ versus control wounds at 2, 4, and 7 dpi. **(C)** Representative flow cytometry scatter plots showing neutrophils (gated on 7-AAD^−^CD45^+^CD11b^+^F4/80^−^Ly6G^+^) in CASP8^iMKO^ and control wounds at 7 dpi. **(D)** Quantification of relative neutrophil frequency (%) in CASP8^iMKO^ versus control (CASP8^fl/fl^) wounds at 7 dpi. **(E)** Representative H&E-stained histological sections of CASP8^iMKO^ and control wounds at 7 dpi. Asterisk indicates hemorrhagic area. Scale bar = 100 μm. **(F)** Quantification of hemorrhagic area (red blood cell area, mm²) within the granulation tissue of CASP8^iMKO^ versus control wounds at 7 dpi. Data are presented as mean ± SEM. Statistical significance was determined by two-tailed Student’s t-test (*p < 0.05; **p < 0.01; ns, not significant). Each dot represents an individual wound. All experiments were performed at least twice independently with a minimum of 5 mice per condition.

### Enhanced macrophage apoptosis preserves tissue architecture despite sustained macrophage loss

To determine whether the tissue defects in FADD^iMKO^ wounds are specific to necroptosis or represent a general consequence of macrophage loss, we analyzed wound repair in cIAP1^iMKO^cIAP2^−/−^ mice, in which combined macrophage-specific deletion of cIAP1 (encoded by *Birc2*) on a germline *Birc3*^−/−^ background enhances macrophage apoptosis. Two features of this model require emphasis. First, the control genotype throughout these experiments is cIAP2^−/−^ rather than wild-type: deletion of *Birc3* alone is not considered sufficient to induce apoptosis, as *Birc2* and *Birc3* are known to have redundant functions (Wong *et al*, 2014). However, *Birc3* deletion was shown to sensitize macrophages to LPS-induced cell death (Conte *et al*, 2006), suggesting that the cIAP2−/− background may represent an apoptosis-sensitized baseline during wound healing.

To determine how macrophage apoptosis impacts tissue repair and architecture, we subjected cIAP1^iMKO^cIAP2^−/−^ mice to excision skin injury. Morphologically, wounds appeared comparable to controls at 7 dpi (**Fig. 2A**). To analyzed immune cell dynamics across the healing time course we subjected single cell suspensions of wound tissue at subsequent phases of repair to flow cytometry analysis. Analysis revealed decreased viable macrophage numbers in cIAP1^iMKO^cIAP2^−/−^ wounds at 2 and 4 dpi (**Fig. 7A, B**). Yet, both cIAP2^−/−^ and cIAP1^iMKO^cIAP2^−/−^ wounds showed reduced macrophage relative numbers when compared to WT wounds (Injarabian *et al*, 2024), suggesting that the cIAP2^−/−^ background could further influence the viability of macrophages in the mid-stage of repair. Analysis of macrophage subpopulations showed reduced Ly6C^high^ macrophages at 2 dpi and reduced Ly6C^low^ reparative macrophages at 4 dpi in cIAP1^iMKO^cIAP2^−/−^ when compared to cIAP2^−/−^ wounds (**Fig. 7C, D**). Together, these findings suggest that both cIAP2^−/−^ and cIAP1^iMKO^cIAP2^−/−^ wounds operate with substantially fewer macrophages in the mid-stage of repair. Consistently, neutrophil numbers were elevated in cIAP1^iMKO^cIAP2^−/−^ wounds at 2 and 4 dpi (**Fig. S9**), and both cIAP2^−/−^ and cIAP1^iMKO^cIAP2^−/−^ wounds showed higher neutrophil numbers at 7 dpi when compared to WT wounds (data not shown).

**Figure 7.**
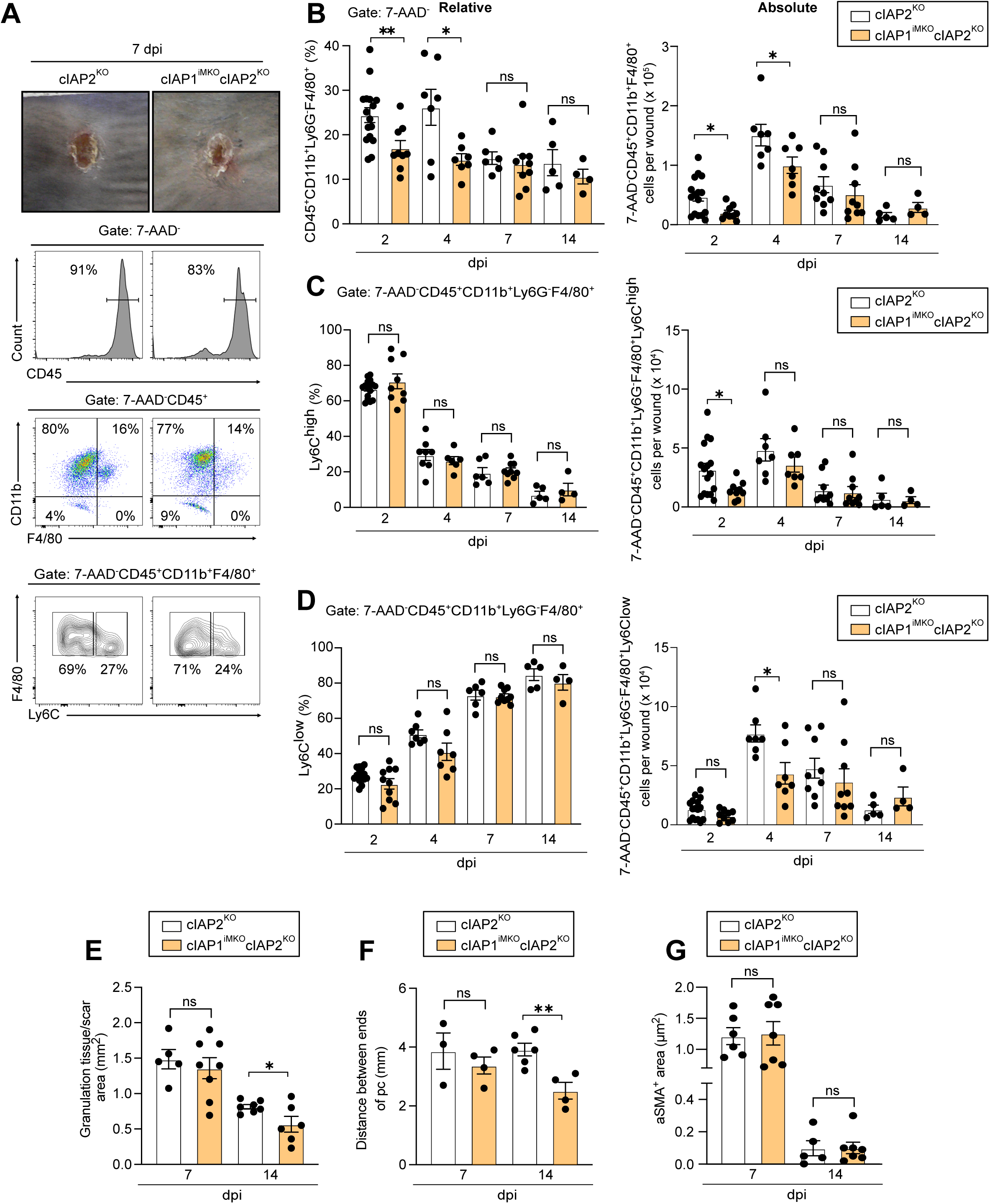
Enhanced macrophage apoptosis reduces macrophage numbers, preserves tissue architecture, and reduces scar formation. **(A)** Representative macroscopic wound images (top) and flow cytometry histograms and scatter plots (bottom) showing viable macrophage gating (7-AAD^−^CD45^+^CD11b^+^Ly6G^−^F4/80^+^) and Ly6C^high^/Ly6C^low^ subpopulation identification in cIAP1^iMKO^cIAP2^KO^ and cIAP2^KO^ control wounds at 7 dpi. **(B)** Quantification of relative (left) and absolute (right) viable macrophage numbers in cIAP1^iMKO^cIAP2^KO^ versus cIAP2^KO^ control wounds across the healing time course (2, 4, 7, and 14 dpi). **(C)** Quantification of relative (left) and absolute (right) Ly6C^high^ macrophage numbers in cIAP1^iMKO^cIAP2^KO^ versus cIAP2^KO^ control wounds across the healing time course. **(D)** Quantification of relative (left) and absolute (right) Ly6C^low^ macrophage numbers in cIAP1^iMKO^cIAP2^KO^ versus cIAP2^KO^ control wounds across the healing time course. **(E)** Quantification of granulation tissue/scar area (mm²) in cIAP1^iMKO^cIAP2^KO^ versus cIAP2^KO^ control wounds at 7 and 14 dpi. **(F)** Quantification of wound contraction, measured as the distance between panniculus carnosus (pc) ends (mm), in cIAP1^iMKO^cIAP2^KO^ versus cIAP2^KO^ control wounds at 7 and 14 dpi. **(G)** Quantification of αSMA^+^ area (µm²) within the granulation tissue of cIAP1^iMKO^cIAP2^KO^ versus cIAP2^KO^ control wounds at 7 and 14 dpi. Data are presented as mean ± SEM. Statistical significance was determined by two-tailed Student’s t-test (*p < 0.05; **p < 0.01; ns, not significant). Each dot represents an individual wound for 7 dpi and 2 pooled wounds for 14 dpi. All experiments were performed at least twice independently with a minimum of 5 mice per condition.

To investigate the consequence of increased macrophage apoptosis on tissue architecture we assessed tissue morphology at 7 and 14 dpi. Analysis revealed normal granulation tissue morphology with no hemorrhagic areas (**Fig. 7E, F; Fig. S9B, C**), demonstrating intact vascular stability and tissue organization despite reduced macrophage numbers. Furthermore, we observed a comparable and normal myofibroblast response (αSMA^+^ area) in cIAP2^−/−^ and cIAP1^iMKO^cIAP2^−/−^ wounds at 7 and 14 dpi (**Fig. 7G**). Despite this comparable myofibroblast response in both conditions, morphometric analysis at 14 dpi revealed reduced scar tissue area and enhanced wound contraction in cIAP1^iMKO^cIAP2^−/−^ wounds when compared to cIAP2^−/−^ controls (**Fig. 7E, F**), suggesting that enhanced macrophage apoptosis might modulate resolution of fibrosis.

Collectively, these results suggest that reducing macrophage numbers through apoptosis is not only more compatible with functional tissue repair, but it could also improve repair quality by reducing scar formation.

## DISCUSSION

Our findings establish a fundamental principle in tissue repair biology: wound healing quality is determined not merely by the presence or absence of macrophages, but by the mode through which macrophages die. By independently manipulating necroptosis and apoptosis using complementary inducible genetic models, we demonstrate that these two RCD modalities exert diametrically opposed effects on tissue architecture, cellular communication networks, and repair quality. Macrophage necroptosis drives structural collapse through selective depletion of reparative IL-4Rα^+^Ly6C^low^ macrophages and disruption of tissue-wide intercellular communication networks, while enhanced macrophage apoptosis preserves structural integrity, sustains a robust reparative fibroblast response, and reduces final scar formation. The phenotypic concordance between FADD^iMKO^ and CASP8^iMKO^ mice firmly attributes the necroptosis-associated defects rather than RCD-independent scaffolding functions of FADD and responsible for repair impairment. Together, these results reframe macrophage cell death as an active, mode-specific determinant of repair outcome, with direct implications for therapeutic targeting in wound disorders and fibrotic disease.

The most consequential cellular phenotype in FADD^iMKO^ wounds was the selective depletion of Ly6C^low^ reparative macrophages at 7 dpi, precisely when these cells are critical for granulation tissue maturation, vascular stabilization, and efferocytosis. Crucially, this depletion occurred without a corresponding increase in Ly6C^low^ macrophage cell death at this stage, implicating impaired phenotypic reprogramming or self-renewal rather than direct cell loss as the underlying mechanism. This interpretation aligns with the established role of IL-4Rα signaling in macrophage proliferative self-renewal during tissue repair (Jenkins *et al*, 2011; Knipper *et al*, 2015), and raises the possibility that DAMPs released by early necroptotic macrophages, including HMGB1, S100 proteins, and extracellular ATP, sustain a pro-inflammatory environment that is incompatible with IL-4-driven phenotypic reprogramming. Notably, HMGB1 released from necrotic cells directly inhibits macrophage efferocytosis through disruption of integrin-dependent engulfment pathways (Friggeri *et al*, 2010), potentially creating a compounding defect in which necroptosis simultaneously limits the generation of reparative macrophages and impairs the efferocytic capacity of those that remain.

The hemorrhagic granulation tissue phenotype of FADD^iMKO^ wounds, phenocopying both a macrophage depletion model (Lucas *et al*, 2010) and myeloid-specific *Il4ra* deletion (Knipper *et al*, 2015) provides compelling evidence that IL-4Rα^+^Ly6C^low^ macrophages are indispensable for vascular integrity beyond their role in vessel formation per se. CD31^+^ vessel density was preserved in FADD^iMKO^ wounds, yet vascular integrity failed. This dissociation between vessel number and vessel function is consistent with the loss of paracrine stabilizing signals, such as angiopoietin-1, PDGF-BB, and TGFβ, normally provided by reparative macrophages to pericytes and endothelial cells (Willenborg *et al*, 2012). Immunofluorescence analysis confirmed a significant reduction in IL-4Rα^+^ macrophages in FADD^iMKO^ wounds at 7 dpi establishing IL-4Rα^+^ macrophage density as a functional correlate of vascular stability in granulation tissue in this model.

The temporal pattern of neutrophil dynamics in FADD^iMKO^ wounds, normal influx at 2 and 4 dpi, significant absolute accumulation at 7 dpi, normalization by 14 dpi, is most consistent with impaired efferocytic clearance driven by low Ly6C^low^ macrophage numbers and reduced efferocytosis efficiency rather than enhanced recruitment. Although MerTK and Axl surface expression on wound macrophages was unchanged when measured by flow cytometry, scRNAseq analysis revealed a broad, cell-type-spanning suppression of eat-me signals alongside divergent efferocytosis receptor transcript changes across fibroblasts, macrophages, dendritic cells and neutrophils, indicating reduced total tissue efferocytic capacity despite normal per-cell receptor expression. Because successful efferocytosis induces anti-inflammatory reprogramming through release of TGFβ, PGE₂, and IL-10 (Bosurgi *et al*, 2017; Fadok *et al*, 1998), we suggest that in our model it could create a self-reinforcing cycle in which reduced efferocytosis lead to impaired neutrophil clearance, sustained DAMP exposure, and continued suppression of the reparative state transition, perpetuating the architectural defects observed throughout the mid-healing phase.

Yet, LIANA+ analysis of scRNAseq data demonstrated that macrophage necroptosis does not simply reduce reparative macrophage numbers but fundamentally remodels the intercellular signaling landscape in wounds. In control wounds, macrophages establish a multivalent stromal-instructive niche coordinating distinct fibroblast subpopulations through three integrated programs: lipid metabolic and scavenger receptor signaling (Apoe–Abca1/Lrp4, Apoc2–Lrp1, Thbs1–/S100a8–/Saa1–Cd36), chemokine-mediated fibroblast priming (Ccl2–Ccr2/Ccr5, Ccl11–Ccr5, Il16–Ccr5), and growth factor–driven remodeling (Wnt5a–Ror1, Gas6–Tyro3, Fgf1/2/7/18–Fgfr4). In FADD^iMKO^ wounds this coordinated program is replaced by a damage-clearance state dominated by hemoglobin-scavenging interactions (Hp–Cd163), complement-associated debris clearance (Gas6–MerTK, Pros1–MerTK, C1qb–C1qbp), and disorganized ECM remodeling cues, a shift that likely contributes to impaired myofibroblast differentiation and architectural collapse beyond what macrophage loss alone would predict. Notably, analogous communication network alterations were already present in unwounded FADD^iMKO^ skin, where macrophage–fibroblast interactions resembled an injury-response state rather than homeostatic tissue maintenance. Consistent with findings in other tissues, necroptosis of Kupffer cells has been shown to remodel the hepatic immune environment by activating neighboring non-immune cells, which in turn drive secondary immune responses (Blériot *et al*, 2015). Moreover, Lyve1⁺ macrophages expand in unwounded FADD^iMKO^ skin in an injury-independent manner and exhibit an interferon-stimulated transcriptional program at 7 dpi, suggesting that specific tissue resident macrophage populations could be sensitive to necroptotic signaling. These findings point to a dual role of macrophage necroptosis, with the potential to drive both inflammatory responses and interferon-associated regenerative programs. DAMPs released by necroptotic cells, including nucleic acids and HMGB1, are well-established activators of pattern recognition receptors such as TLR3 and the cGAS–STING pathway, which induce type I interferon production. Interferon signaling has, in turn, been linked to tissue regeneration: TLR3-mediated sensing of endogenous dsRNA is required for wound-induced hair follicle neogenesis in skin (Nelson *et al*, 2015), and Lyve1^+^ macrophages have recently been associated with neonatal heart regeneration (Chapman *et al*, 2025). The transcriptional state of Lyve1⁺ macrophages in FADD^iMKO^ skin may therefore reflect activation downstream of necroptotic danger signals, pointing to a regenerative program. Notably, this response appears spatially uncoupled from the wound, as these cells expand predominantly in the peri-wound dermis rather than within the granulation tissue.

Together, these observations support the emerging concept that necroptotic outcomes are highly context-dependent, shaped by the identity of the dying cell, the inflammatory state of the tissue, and the timing of the death signal (Zhou *et al*, 2020).

In striking contrast, increased macrophage apoptosis in cIAP1^iMKO^cIAP2^−/−^ mice preserved fundamental tissue architecture and even improved repair quality despite operating with substantially fewer macrophages throughout healing.

A critical observation within the cIAP1^iMKO^cIAP2^−/−^ model further challenges conventional interpretations of how inflammatory resolution relates to repair quality. Neutrophil numbers remained elevated at 2 and 4 dpi in cIAP1^iMKO^cIAP2^−/−^ wounds relative to cIAP2^−/−^ controls, and elevated in both cIAP2^−/−^ and cIAP1^iMKO^cIAP2^−/−^ wounds when compared to WT wounds at 7 dpi. Yet, despite this delayed inflammatory resolution, no significant differences in tissue architecture could be observed at 7 dpi when compared to WT wounds: no hemorrhagic areas, intact granulation tissue morphology and unimpaired myofibroblast differentiation. This stands in sharp contrast to FADD^iMKO^ wounds, where comparable neutrophil accumulation was accompanied by vasculature collapse. The dissociation between neutrophil persistence and tissue outcome in the cIAP model implies that the degree of inflammatory resolution, measured as neutrophil clearance kinetics, is not the primary determinant of repair quality. Rather, it is the mode of macrophage death, and the downstream tissue signaling environment it creates, that shapes the repair trajectory. In FADD^iMKO^ wounds, neutrophil accumulation occurs within a necroptotic DAMP-rich milieu that disrupts vascular stability, impairs macrophage state transitions, and collapses the reparative stromal-instructive communication program. In cIAP1^iMKO^cIAP2^−/−^ wounds, neutrophil persistence occurs in a tissue context where membrane integrity is preserved, DAMPs would be are absent, and efferocytic reprogramming of surviving macrophages sustains reparative signaling. These data suggest that repair quality is governed by mechanisms that operate largely independently of the conventional pro-versus anti-inflammatory axis, specifically, by the qualitative character of cell death and the paracrine programs it triggers in phagocytes, rather than by the speed or completeness of inflammatory cell clearance. This reinterpretation has direct consequences for understanding the myofibroblast response in cIAP1^iMKO^cIAP2^−/−^ wounds. Despite a reduced macrophage pool, the αSMA^+^ myofibroblast area at 7 dpi was quantitatively intact, representing a proportionally augmented fibrotic response per remaining macrophage when considered against the control references (Injarabian *et al*, 2024; Willenborg *et al*, 2012). We propose that efferocytosis of the elevated apoptotic macrophage burden by surviving phagocytes could constitute an active instructive signal that compensates for the numerical deficit. Indeed, it was demonstrated in the context of human macrophages that exposing macrophages to apoptotic neutrophils prior to co-culturing with fibroblasts increases their pro-fibrotic potential (Maksimova *et al*, 2023). Mechanistically, TGFβ and IL-10, both upregulated during efferocytic reprogramming (Bosurgi *et al*, 2017; Fadok *et al*, 1998) could promote fibroblast-to-myofibroblast differentiation. Bosurgi et al. further established that apoptotic cell recognition is a necessary co-stimulus for IL-4/IL-13-driven macrophage tissue repair gene expression (Bosurgi *et al*, 2017). Therefore, we hypothesize that in cIAP1^iMKO^cIAP2^−/−^ wounds, the elevated apoptotic burden could provide precisely this co-stimulus to macrophages operating in an IL-4-enriched wound environment, potentially amplifying reparative programming per surviving cell and sustaining robust fibroblast activation despite reduced macrophage numbers.

Overall, our findings suggest that the mode of cell death is a previously underappreciated determinant of macrophage function, one that shapes the trajectory of tissue repair through mechanisms operating largely outside the conventional framework of pro-versus anti-inflammatory balance.

## MATERIALS AND METHODS

### Animals

Mouse strains are listed in Table 1. Procedures were authorized by LANUV (81-02.04.2020.A466). Mice were housed 2-5 per cage at 22-24°C, 45-65% humidity, 12h light/dark cycle, with standard food (Altromin) and water ad libitum. Hygiene monitoring followed FELASA recommendations (SPF status). FADDiMKO mice were generated by crossing Faddfl/fl (Mc Guire, 2010) with Cx3cr1CreER (Yona, 2013) (C57BL/6N). cIAP1iMKOcIAP2-/- mice were generated by crossing Birc2fl/flBirc3-/- (Wong et al. 2014) with Cx3cr1CreER. CASP8iMKO mice were generated by crossing Casp8fl/fl with Cx3cr1CreER. Cx3Cr1YFP reporter mice (Yona, 2013) were used for validation. All mice were generated by in-house mating. Male or female mice aged 8-20 weeks were used. Tamoxifen (3 mg in 0.1 ml corn oil) was injected i.p. daily for 5 consecutive days before wounding. Genotyping primers are in Table 2.

**Table 1.** Mouse strains.

| Experimental models (strain) | Reference | Link |
| --- | --- | --- |
| C57BL/6J mice | Charles River | https://www.criver.com/products-services/find-model/jax-c57bl6j-mice?region=23 |
| *Fadd*<sup>fl/fl</sup> mice | (Mc Guire *et al*, 2010) | N/A |
| *Cx3cr1*-creER mice | (Yona *et al*, 2013) | N/A |
| *Fadd*<sup>fl/fl</sup> *Cx3cr1*<sup>CreER</sup> mice | (generated in this study) | N/A |
| *Cx3Cr1*<sup>ER-YFP</sup> mice | (Yona *et al*, 2013) | N/A |
| *Caspase8*<sup>fl/fl</sup> mice | (Beisner *et al*, 2005) | N/A |
| *Caspase8*<sup>fl/fl</sup> *Cx3cr1*<sup>CreER</sup> mice | (generated in this study) | N/A |
| *Birc2*<sup>fl/fl</sup> *Birc3*<sup>-/-</sup> mice | (Wong *et al*, 2014) | N/A |
| *Birc2*<sup>fl/fl</sup> *Birc3*<sup>-/-</sup> *Cx3cr1*<sup>CreER</sup> mice | (generated in this study) | N/A |

**Table 2.**
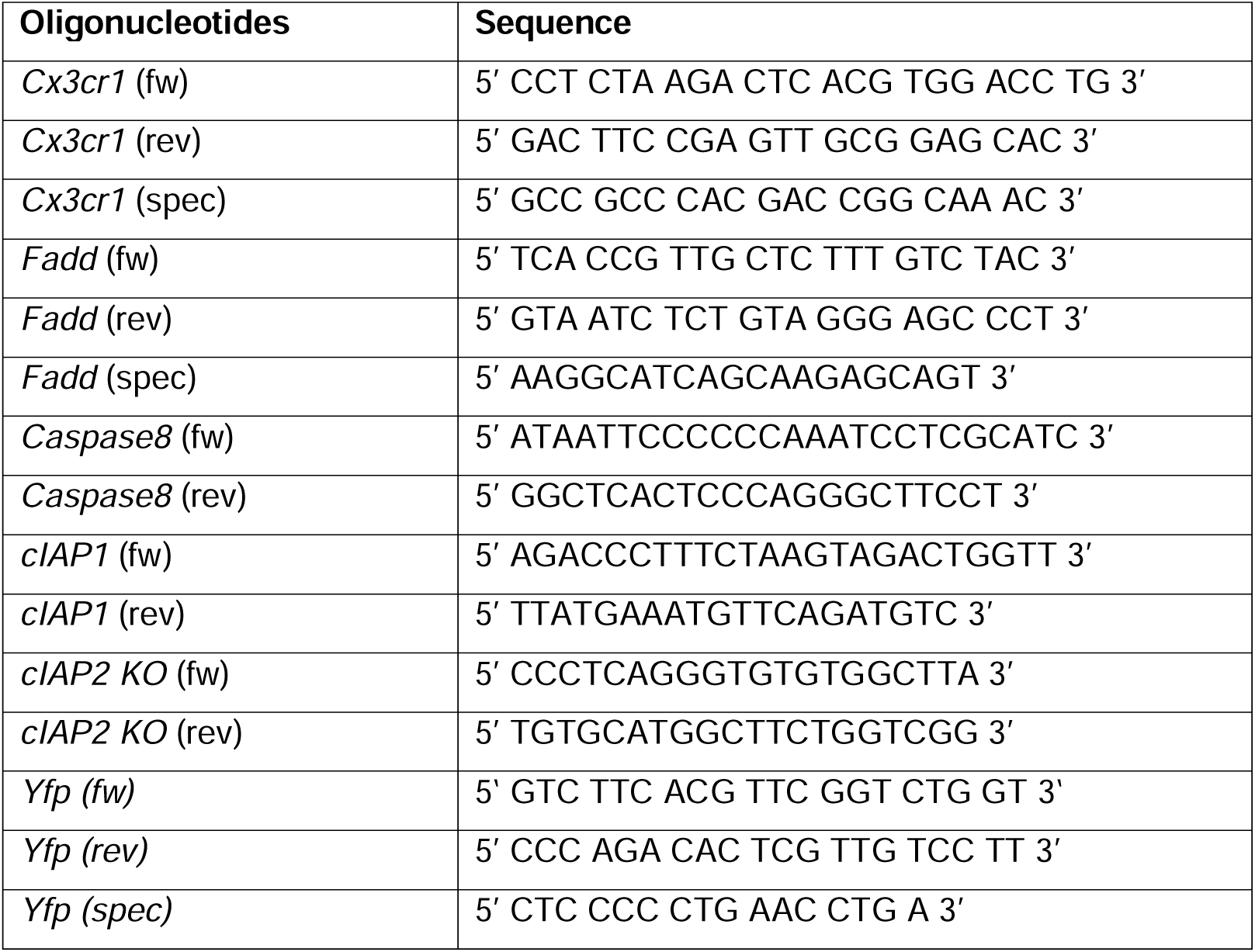
Primer sequences for genotyping.

### Excisional punch injury

Mice were anesthetized (Ketavet 100 mg/kg + Rompun 10 mg/kg, i.p.), back skin shaved and disinfected with 70% ethanol. Full-thickness 6 mm punch biopsies were created. Wounds were excised at indicated timepoints, bisected caudocranially, and either fixed in Roti®Histofix or embedded in O.C.T. compound and stored at −80°C (Injarabian *et al*, 2024; Knipper *et al*, 2015; Lucas *et al*, 2010; Willenborg *et al*, 2012).

### Morphometric analysis

H&E-stained paraffin sections were analyzed using light microscopy (Leica DM4000B) with KY-F75U digital camera (JVC) and Diskus 4.50 software. Granulation tissue area, epithelial gap, and hemorrhagic area were quantified. Analysis was performed blinded.

### Immunostaining

Cryosections (10μm) were fixed in acetone (−20°C) or 4% PFA, blocked (0.1% Triton/1% BSA/10% normal goat serum in PBS, 1h RT), then incubated with primary antibodies (2h RT or overnight 4°C) in antibody diluent (DCS). After washing, sections were incubated with fluorophore-conjugated secondary antibodies (1h RT), washed, and mounted in glycerol gelatine. Primary antibodies: anti-αSMA-Cy3 (Sigma C6198, 1:250), anti-F4/80 (Dianova BMA, 1:200), anti-CD31 (1:200), anti-Lyve1 (Abcam ab14917, 1:50), anti-IL-4Rα (Santa Cruz sc-28361, 1:200), anti-Ly6G. Secondary antibodies: Goat anti-rat Alexa Fluor 488 (Thermo Fisher A-11006), Goat anti-rabbit Alexa Fluor 594 (Thermo Fisher A-11012). DNA staining: DAPI (Sigma, 1 µg/mL). Images acquired with Keyence BZ-9000 (10× or 20× objective) or Leica TCS-SP8 confocal microscope (20× or 40× objective). Analysis used ImageJ.

### Flow cytometry

Single-cell suspensions were prepared by enzymatic digestion (Liberase TM, 30 mg/mL, Roche, 90 min 37°C shaking) and mechanical disruption (Medimachine, BD Biosciences, 5 min). Cells were filtered (70 μm, 40 μm) and washed (PBS/1% BSA/2 mM EDTA). Fc receptors were blocked (anti-CD16/CD32, 1:50, Thermo Fisher). Surface staining: FITC or Pacific blue anti-CD45 (1:200), APC or APC-Cy7 anti-CD11b (1:300), PE anti-F4/80 (1:50, Bio-Rad), PE-Cy7 Ly6C (1:400, BD), APC-Cy7 Ly6G (1:400, BD). Apoptosis: Pacific blue Annexin V (Thermo Fisher) for 25 min RT in Annexin V binding buffer. Efferocytosis markers: PE-Cy7 anti-MerTK (1:50), APC anti-Axl (1:100), Pacific blue anti-Lyve1 (1:200). Dead cells excluded: 7-AAD (Thermo Fisher). Analysis: FACSCanto II (BD). Data analyzed: FlowJo 10.8.2.

### Single-cell RNA sequencing and LIANA analysis

Single-cell suspensions of wound tissue from FADD^iMKO^ and control mice were prepared at day 0 (unwounded skin), 2 dpi, and 7 dpi as described in Willenborg et al., 2021 (Willenborg *et al*, 2021). For unwounded skin, 8 6-mm punches of back skin were used per mice, 2 mice per genotype. For each injury timepoint and genotype, 5 wounds from 3 mice were pooled to obtain sufficient cell numbers. Cells were resuspended in PBS supplemented with 0.04% bovine serum albumin at a concentration of 1,000 cells/µl. Single-cell capture, barcoding, and library preparation were performed using the 10x Genomics Chromium platform (Chromium Next GEM Single Cell 3′ Reagent Kit v3.1) according to the manufacturer’s instructions, targeting a recovery of 10,000-20,000 cells per sample. Libraries were sequenced on an Illumina NovaSeq 6000 instrument using a PE100 protocol, aiming at a minimum sequencing depth of 50,000 reads per cell.

LIANA+ analysis was performed according to Dimitrov et al. 2024. Only cell populations with ≥30 cells (mc30) were included in the LIANA+ analysis. Statistics

Student’s unpaired two-tailed t-test (2 groups) or one-way ANOVA with Tukey’s post-test (>2 groups) using GraphPad Prism 9. Welch correction applied for unequal variances. P < 0.05 considered significant. All P values reported in figure legends.

## DATA AVAILABILITY STATEMENT

Datasets available at [GEO accession number to be added]

## ACKNOWLEDGEMENTS

This work was supported by Deutsche Forschungsgemeinschaft (DFG): CRC 1403 (414786233), Excellence Strategy CECAD (EXC 2030-390661388), and Center for Molecular Medicine Cologne.

## AUTHOR CONTRIBUTIONS

Conceptualization: LI, SW, SAE; Investigation: LI, SW, DW, NR, YB, JT, ES; Methodology: LI, SW, DES, JT; Resources: MP, HK; Visualization: LI, NR; Writing-original draft: LI, SAE; Writing-review & editing: All authors; Funding Acquisition: SAE; Supervision: LI, SAE.

## SUPPLEMENTARY FIGURE LEGENDS

**Supplementary Figure S1 related to Figure 1. Validation of tamoxifen induction protocol and cell death kinetics in FADD**^iMKO^ **and cIAP1**^iMKO^**cIAP2**^KO^ **models. (A)** Experimental timeline showing tamoxifen administration (3 mg/ml, intraperitoneally, 5 consecutive days), wounding at day 0, and analysis timepoints (2, 4, and 7 dpi). **(B)** Quantification of 7-AAD^+^ macrophage frequency (%) gated on CD45^+^CD11b^+^Ly6G^−^F4/80^+^ cells in FADD^iMKO^ versus control wounds at 4, 7, and 14 dpi. **(C)** Quantification of 7-AAD^+^ cells within the Ly6C^high^ macrophage subpopulation (CD45^+^CD11b^+^Ly6G^−^F4/80^+^Ly6C^high^) in FADD^iMKO^ versus control wounds at 2, 4, and 7 dpi. **(D)** Quantification of 7-AAD^+^ cells within the Ly6C^low^ macrophage subpopulation (CD45^+^CD11b^+^Ly6G^−^F4/80^+^Ly6C^low^) in FADD^iMKO^ versus control wounds at 2, 4, and 7 dpi. **(E)** Quantification of Annexin-V^+^ cells within the Ly6C^high^ macrophage subpopulation (7-AAD^−^CD45^+^CD11b^+^F4/80^+^Ly6C^high^) in cIAP1^iMKO^cIAP2^KO^ versus cIAP2^KO^ control wounds at 2 dpi. **(F)** Quantification of Annexin-V^+^ cells within the Ly6C^low^ macrophage subpopulation (7-AAD^−^CD45^+^CD11b^+^F4/80^+^Ly6C^low^) in cIAP1^iMKO^cIAP2^KO^ versus cIAP2^KO^ control wounds at 2 dpi. Data are presented as mean ± SEM. Statistical significance was determined by two-tailed Student’s t-test (*p < 0.05; **p < 0.01; ns, not significant). Each dot represents an individual wound. All experiments were performed at least twice independently with a minimum of 5 mice per condition.

**Supplementary Figure S2 related to Figure 2. Tissue morphology and IL-4R**α^+^ **macrophage analysis in FADD**^iMKO^ **wounds. (A)** Quantification of wound contraction, measured as the distance between panniculus carnosus (pc) ends (mm), in FADD^iMKO^ versus control wounds at 7 dpi. **(B)** Quantification of total granulation tissue area (mm²) in FADD^iMKO^ versus control wounds at 7 dpi. **(C)** Quantification of CD31^+^ endothelial area (mm²) within the granulation tissue of FADD^iMKO^ versus control wounds at 7 dpi. **(D)** Schematic diagram of wound tissue anatomy (left) indicating the imaging region (red box). Representative immunofluorescence images (right) of wound granulation tissue from control and FADD^iMKO^ mice at 7 dpi co-stained with anti-F4/80 (red), anti-IL-4Rα (green), and DAPI (blue). Arrows indicate F4/80^+^IL-4Rα^+^ double-positive macrophages. Dotted lines delineate the epithelium. he, hyperproliferative epithelium; g.t., granulation tissue; dWAT, dermal white adipose tissue; pc, panniculus carnosus. Scale bar = 50 μm. **(E)** Quantification of IL-4Rα^+^F4/80^+^ macrophages per high-power field (HPF) in the granulation tissue of FADD^iMKO^ versus control wounds at 7 dpi. Data are presented as mean ± SEM. Statistical significance was determined by two-tailed Student’s t-test (*p < 0.05; ns, not significant). Each dot represents an individual wound. All experiments were performed at least twice independently with a minimum of 5 mice per condition.

**Supplementary Figure S3 related to Figure 3. Increased neutrophil density in FADD**^iMKO^ **wounds. (A)** Schematic diagram of wound tissue anatomy (left) indicating the imaging region (red box). Representative immunofluorescence images (right) of wound granulation tissue from control and FADD^iMKO^ mice at 7 dpi stained with anti-Ly6G (red; neutrophil marker) and DAPI (blue). Arrows indicate Ly6G^+^ neutrophils. g.t., granulation tissue; dWAT, dermal white adipose tissue; pc, panniculus carnosus. Scale bar = 20 μm. **(B)** Quantification of Ly6G^+^ cells per high-power field (HPF) in the granulation tissue of FADD^iMKO^ versus control wounds at 7 dpi. Data are presented as mean ± SEM. Statistical significance was determined by two-tailed Student’s t-test (**p < 0.01). Each dot represents an individual wound. All experiments were performed at least twice independently with a minimum of 5 mice per condition.

**Supplementary Figure S4 related to Figure 3. scRNAseq cell type annotation strategy. (A)** UMAP visualization of the integrated scRNAseq dataset from unwounded skin (day 0), 2 dpi, and 7 dpi samples, colored by timepoint. **(B)** UMAP visualization of cell clusters from the 7 dpi dataset colored by annotated cell type, with major populations outlined and labeled: Fibroblast (FB), DC, Macrophage (M), Neutrophil (N). Cell types used as input for LIANA+ ligand–receptor interaction analysis are indicated.

**Supplementary Figure S5 related to Figure 4. Unsupervised cluster annotation for scRNAseq dataset used in LIANA+ analysis. (A)** UMAP visualization at clustering resolution 0.4, with 23 clusters (0–22) annotated by color. **(B)** Bar plots showing the top marker gene expression profiles for each cluster, used for cell-type annotation. **(C)** UMAP visualizations of each individual sample split by genotype and timepoint: D0_NH_neg (unwounded skin, Ctrl), D0_NH_pos (unwounded skin, FADD^iMKO^), D2_neg (2 dpi, Ctrl), D2_pos (2 dpi, FADD^iMKO^), D7_neg (7 dpi, Ctrl), and D7_pos (7 dpi, FADD^iMKO^), colored by cluster identity at resolution 0.4.

**Supplementary Figure S6 related to Figure 4. Reduced myofibroblast accumulation in FADD**^iMKO^ **wounds assessed by scRNAseq. (A)** Line graph showing the percentage of myofibroblasts among total sequenced cells in FADD^iMKO^ and control wounds at day 0 (unwounded skin), 2 dpi, and 7 dpi, as inferred from scRNAseq cell-type annotation. Each timepoint represents data from pooled samples of the indicated genotype.

**Supplementary Figure S7 related to Figure 4. Characterization of Lyve1**^+^ **macrophage expansion in FADD**^iMKO^ **mice. (A)** Table showing absolute numbers and percentages of Lyve1^+^ macrophages from total macrophages in control and FADD^iMKO^ wounds at 2 and 7 dpi, as inferred from scRNAseq data. **(B)** Volcano plot of differentially expressed genes in Lyve1^+^ macrophages from FADD^iMKO^ versus control wounds at 7 dpi (adjusted p < 0.05). Selected genes of interest are labeled. **(C)** Table listing top differentially expressed genes in Lyve1^+^ macrophages from FADD^iMKO^ versus control wounds at 7 dpi, including adjusted p-value (Padj) and log2 fold-change (Log2FC). Highlighted genes: *Egfr* (upregulated; macrophage proliferation) and *Cd36* (downregulated; efferocytosis receptor). **(D)** Gene Ontology (GO) enrichment dot plot showing biological processes enriched among genes upregulated in Lyve1^+^ macrophages from FADD^iMKO^ versus control wounds 7 dpi. Dot size indicates gene count; color indicates adjusted p-value. Enriched terms include antiviral defense, regulation of innate immune response, and interferon-response pathways. **(E)** Feature plots showing *Lyve1* expression in the macrophage cluster from scRNAseq of unwounded skin from control and FADD^iMKO^ mice, demonstrating pre-injury expansion of Lyve1^+^ macrophages in FADD^iMKO^ skin.

**Supplementary Figure S8 related to Figure 5. Altered macrophage–fibroblast communication in FADD**^iMKO^ **unwounded skin. (A)** Dot plot showing unique ligand–receptor interactions specific to control unwounded skin, inferred by LIANA+ analysis. **(B)** Dot plot showing unique ligand–receptor interactions specific to FADD^iMKO^ unwounded skin. Dot color indicates expression magnitude; dot size indicates interaction specificity.

**Supplementary Figure S9 related to Figure 7. Neutrophil dynamics and tissue morphology in cIAP1**^iMKO^**cIAP2**^KO^ **wounds. (A)** Quantification of relative (left) and absolute (right) neutrophil numbers (gated on 7-AAD^−^CD45^+^CD11b^+^F4/80^−^Ly6G^+^) in cIAP1^iMKO^cIAP2^KO^ versus cIAP2^KO^ control wounds across the healing time course (2, 4, 7, and 14 dpi). **(B)** Representative H&E-stained histological sections of cIAP2^KO^ control (top) and cIAP1^iMKO^cIAP2^KO^ (bottom) wounds at 7 dpi showing normal granulation tissue formation in both conditions and absence of hemorrhagic areas in cIAP1^iMKO^cIAP2^KO^ wounds. Scale bar = 200 μm. **(C)** Quantification of hemorrhagic area (red blood cell area, mm²) within the granulation tissue of cIAP1^iMKO^cIAP2^KO^ versus cIAP2^KO^ control wounds at 7 dpi. Data are presented as mean ± SEM. Statistical significance was determined by two-tailed Student’s t-test (ns, not significant). Each dot represents an individual wound for 7 dpi and 2 pooled wounds for 14 dpi. All experiments were performed at least twice independently with a minimum of 3 mice per condition.

