## Supplementary figures and images for "Modes of programmed macrophage cell death govern outcome of cutaneous wound healing"

### Supplementary figure 1

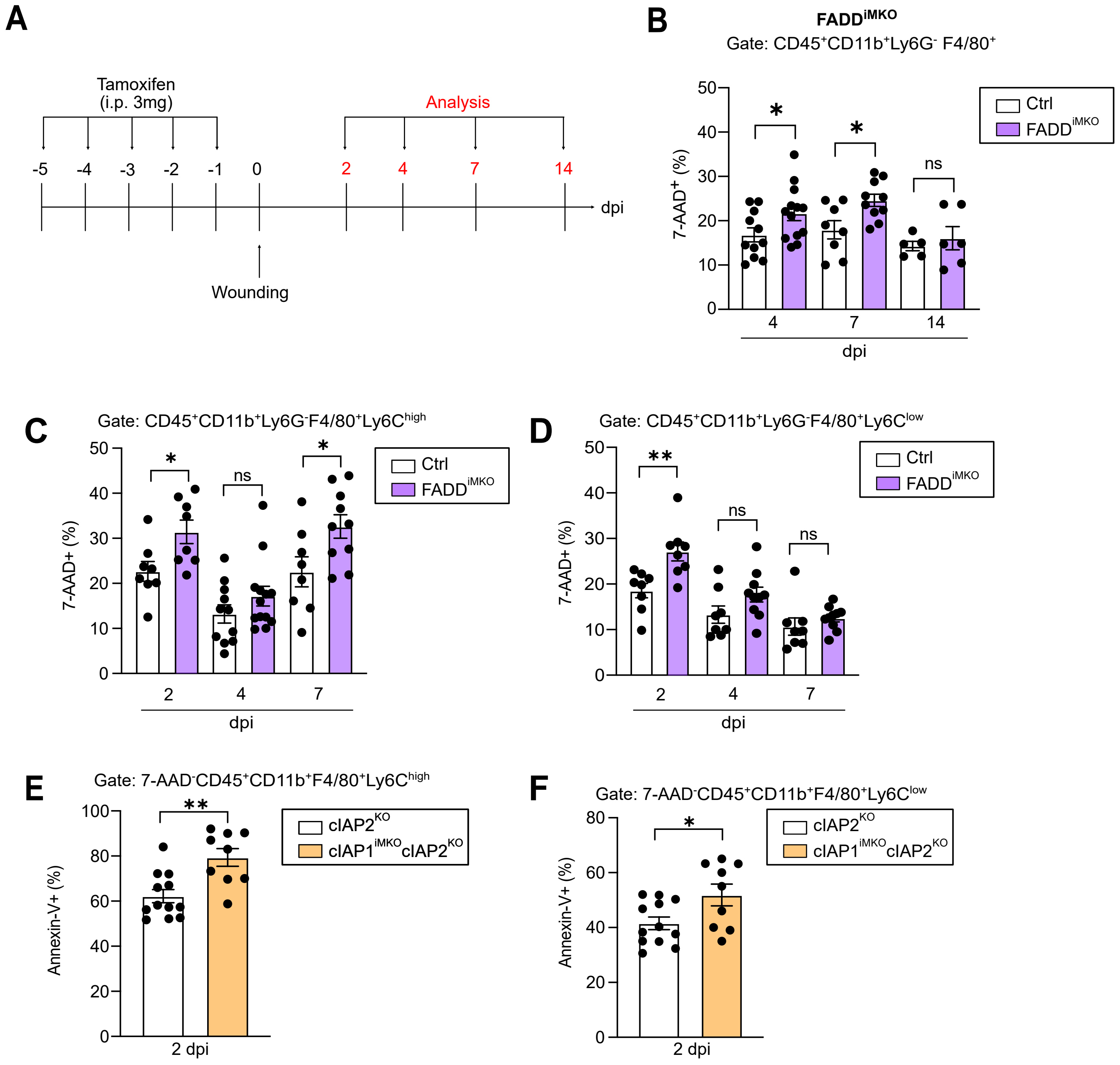

### Supplementary figure 2

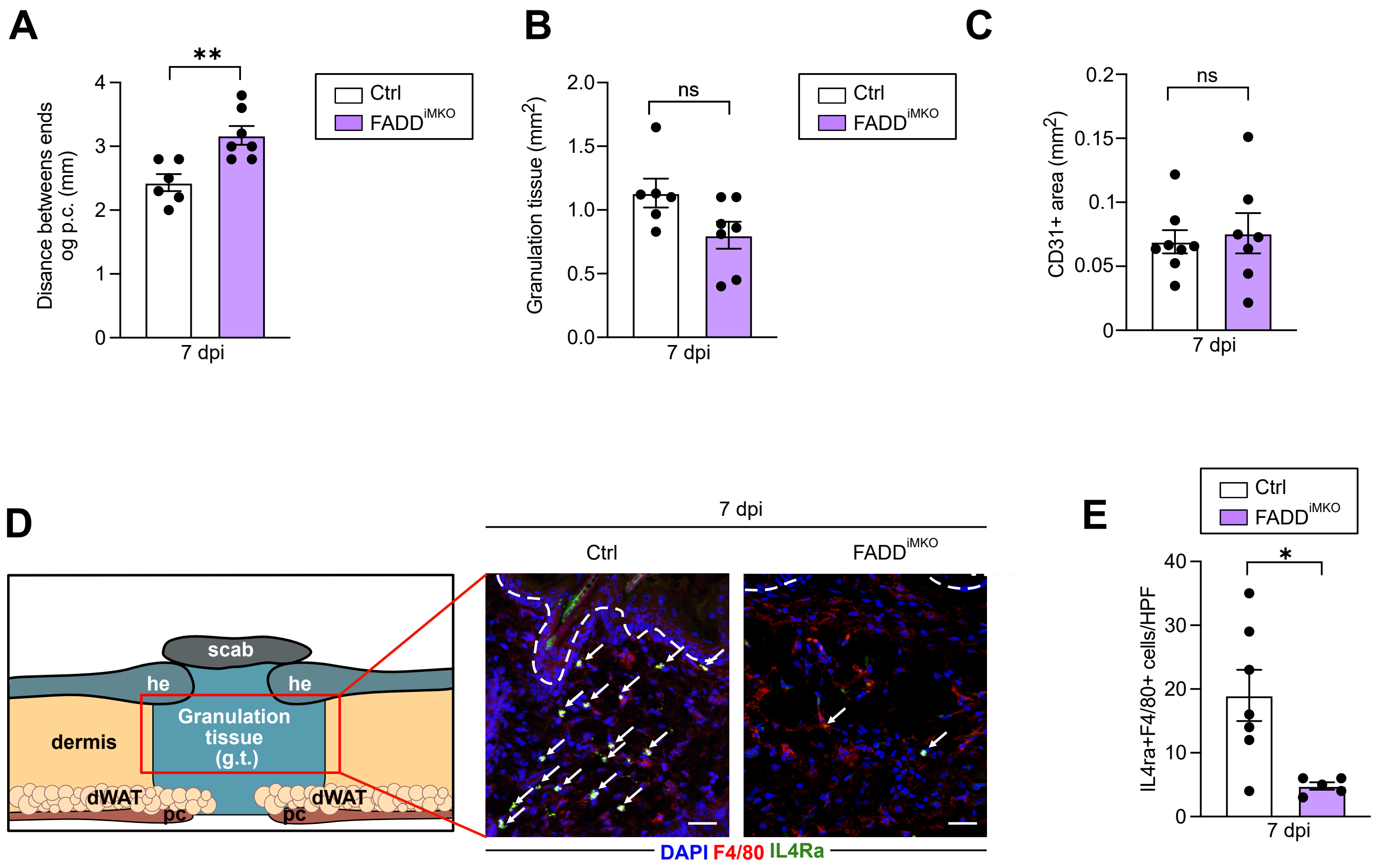

### Supplementary figure 3

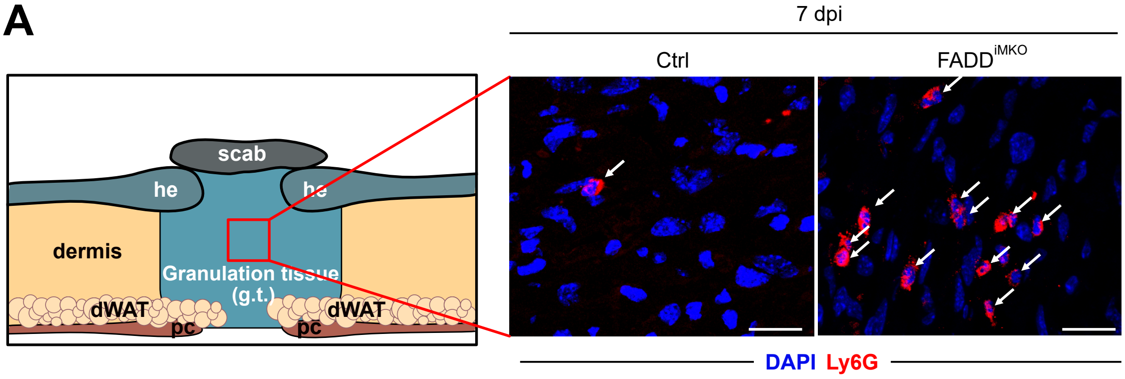

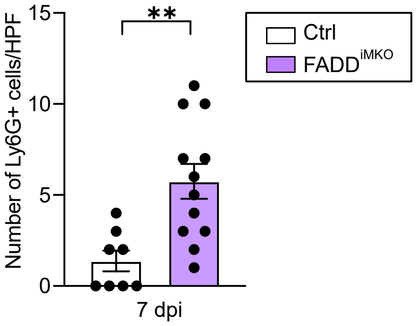

### Supplementary figure 4

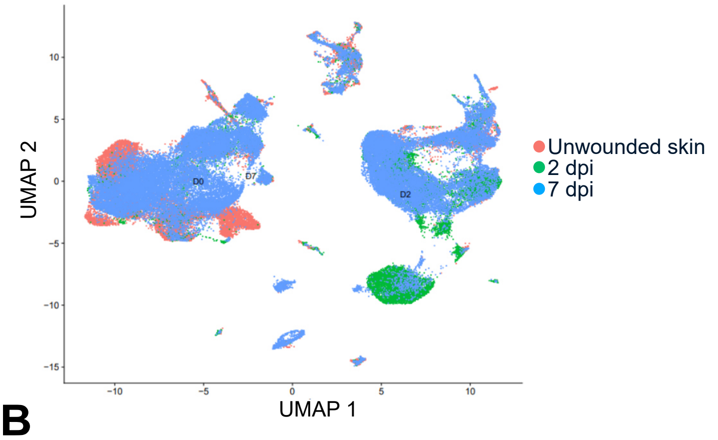

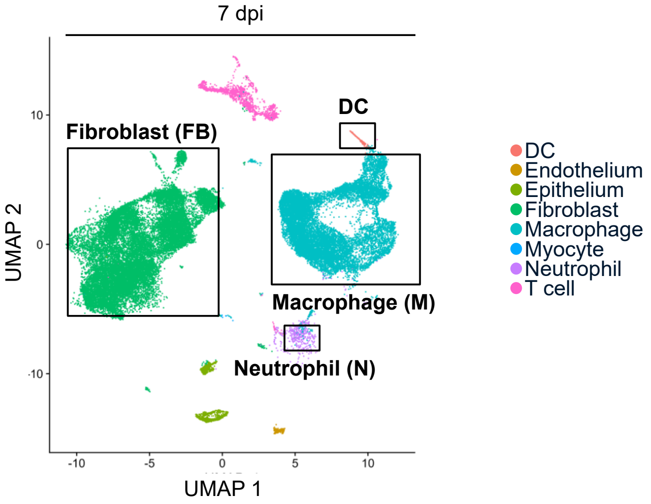

### Supplementary figure 5

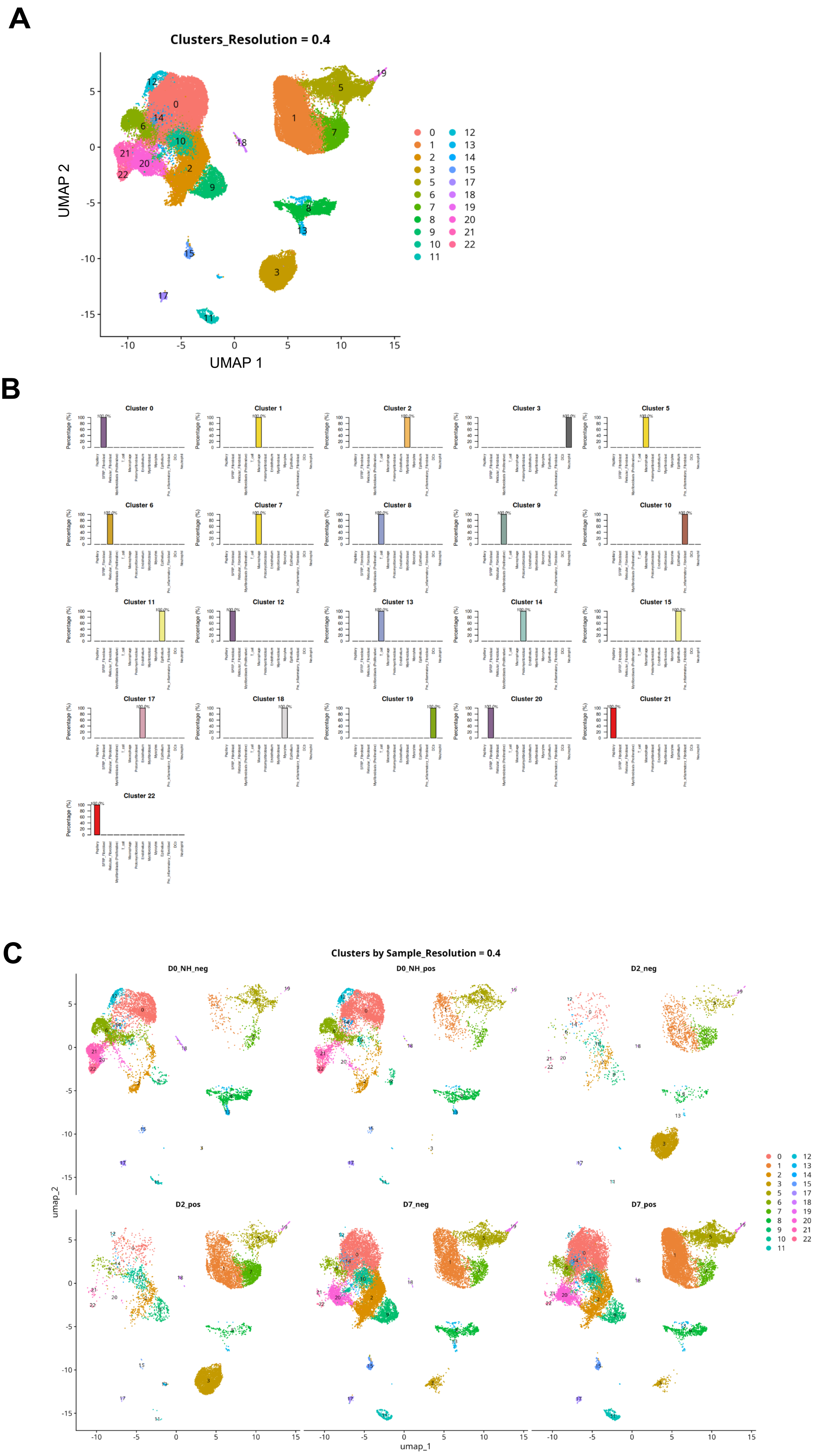

### Supplementary figure 6

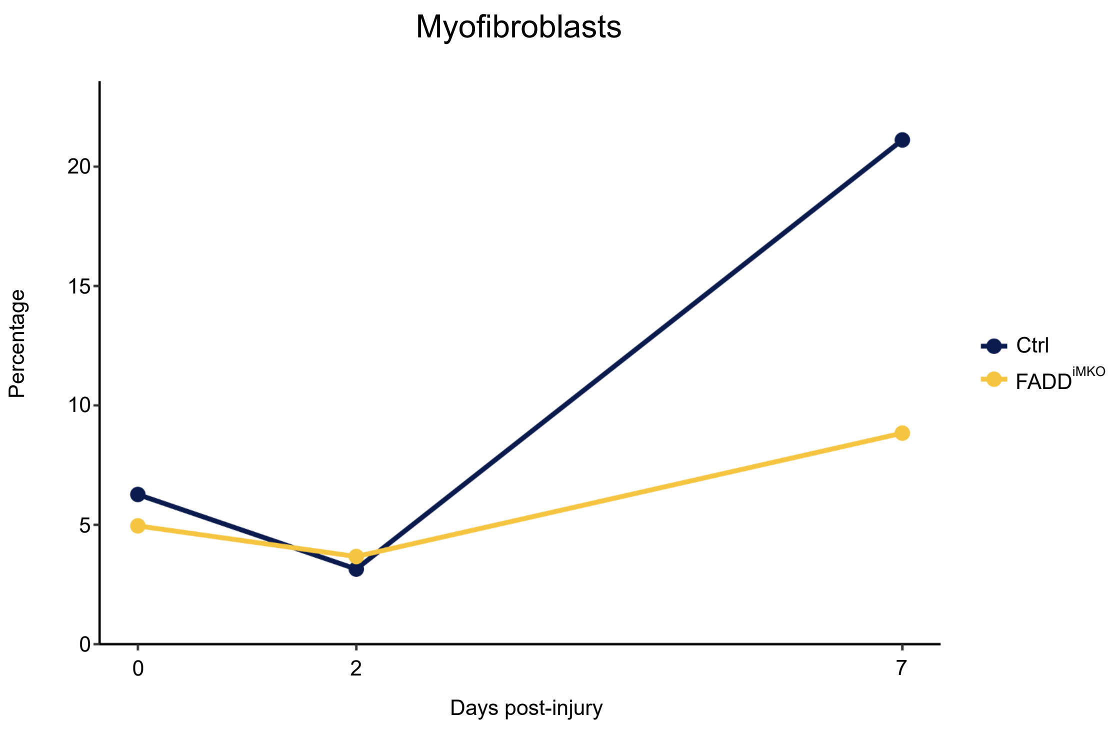

### Supplementary figure 8

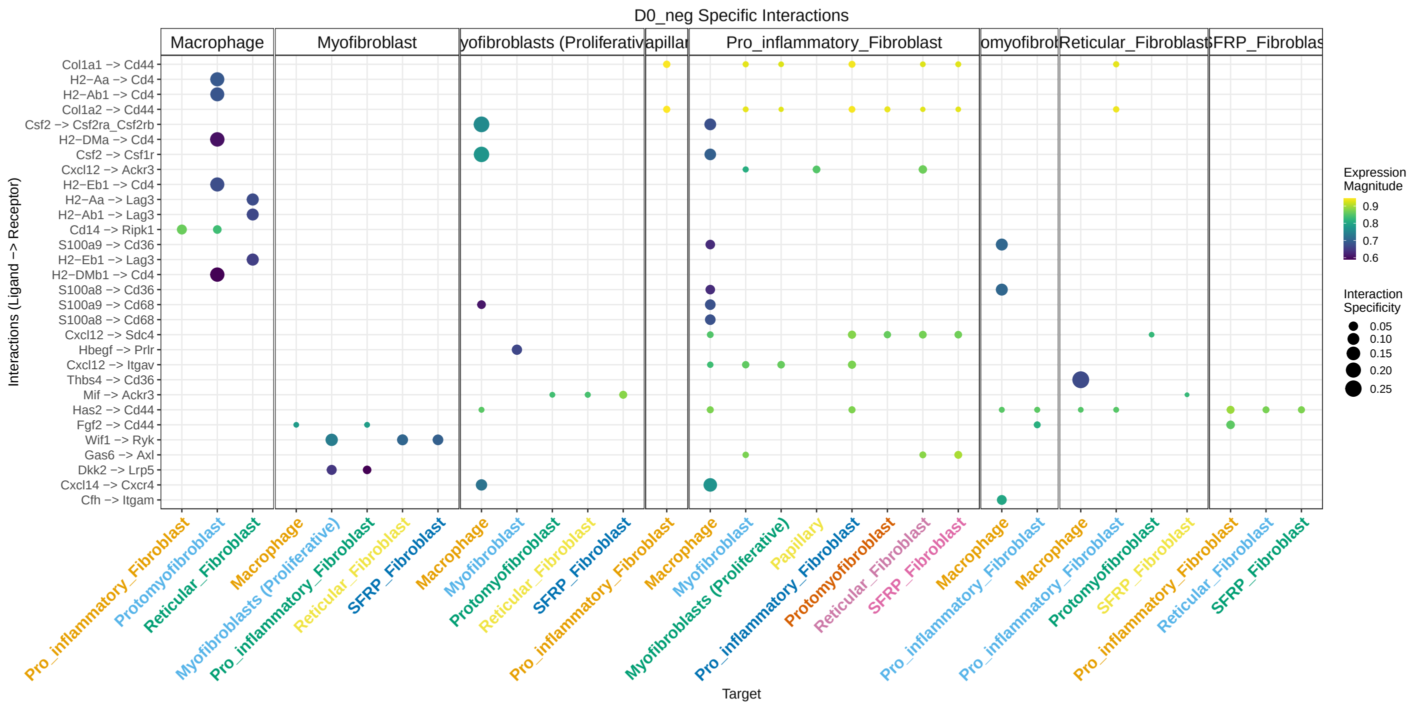

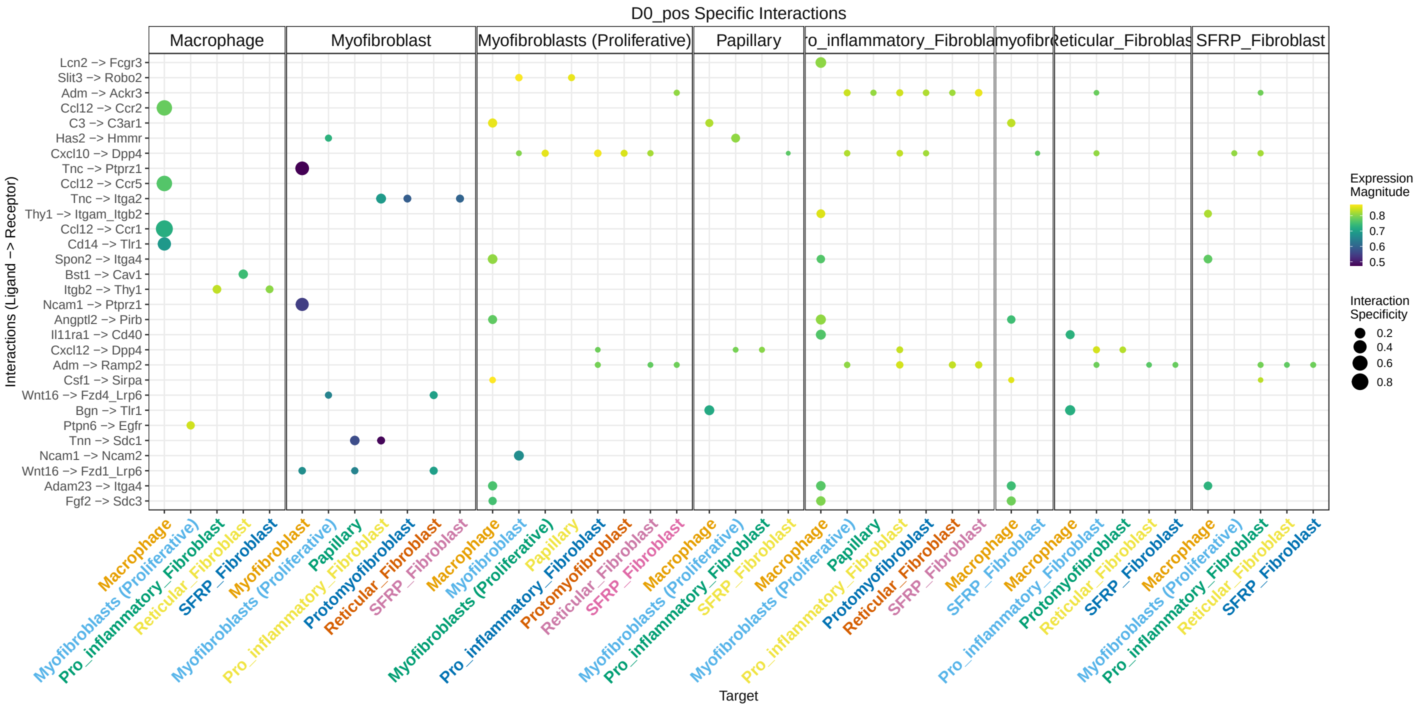

### Supplementary figure 9

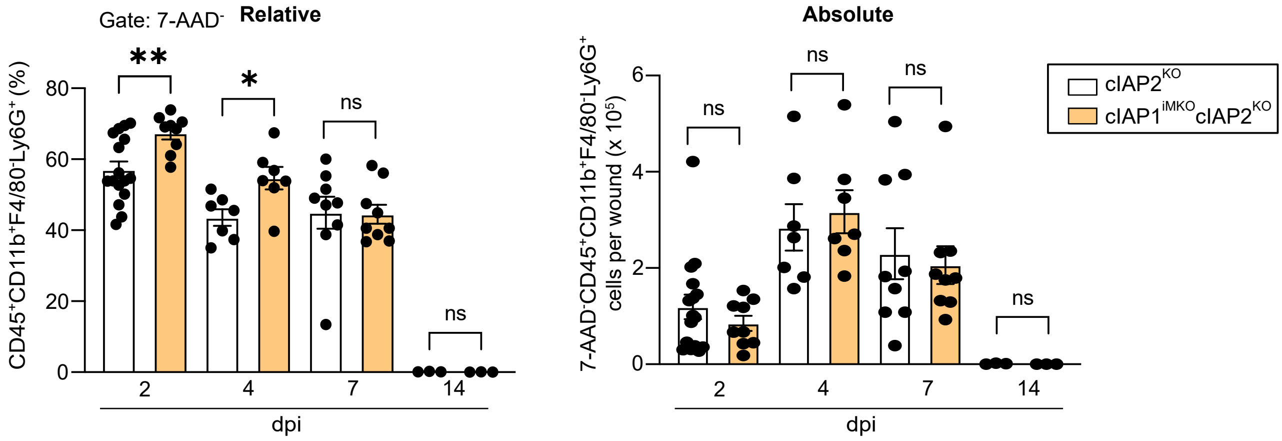

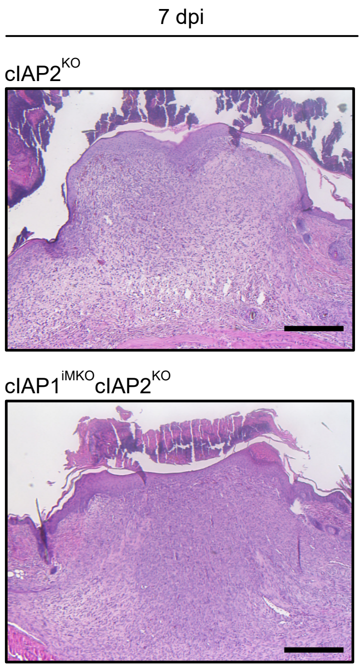

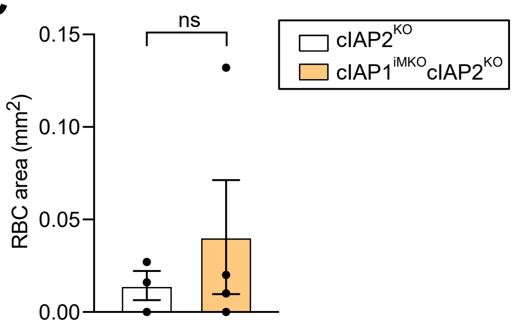
