## Supplementary figure 7 for "Modes of programmed macrophage cell death govern outcome of cutaneous wound healing"

**A**

| Cell-type | Genotype | Timepoint | N (Lyve1+ macrophages) | N (all macrophages) | Percentage [%] of Lyve1+ macrophages from total macrophages |
|---|---|---|---|---|---|
| Macrophages | Control | 2 dpi | 3 | 1277 | 0.235 |
| Macrophages | FADDiMKO | 2 dpi | 27 | 4724 | 0.572 |
| Macrophages | Control | 7 dpi | 225 | 4919 | 4.574 |
| Macrophages | FADDiMKO | 7 dpi | 1225 | 9072 | 13.503 |

**B**

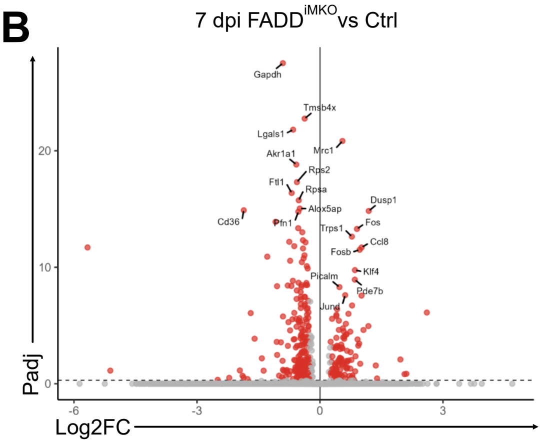


**C**

| Gene name | Padj | Log2FC |
|---|---|---|
| Slc15a2 | 3.12E-11 | 2.60646622 |
| Gm26532 | 5.65E-06 | 2.10768386 |
| Auts2 | 6.07E-06 | 2.06241134 |
| **Egfr** | **3.35E-07** | **1.96136783** |
| Clec3b | 1.42E-05 | 1.39262631 |
| Pi16 | 1.10E-06 | 1.360651 |
| Dusp1 | 5.97E-20 | 1.18946905 |
| Pde4d | 4.57E-10 | 1.1809811 |
| Serping1 | 2.67E-05 | 1.11698839 |
| Angptl4 | 3.50E-11 | -1.6897269 |
| Gt(ROSA)26Sor | 1.60E-05 | -1.7653175 |
| **Cd36** | **5.21E-20** | **-1.8581717** |
| Qrsl1 | 1.19E-05 | -1.8623272 |
| Gm47428 | 8.34E-06 | -1.888034 |
| Gm32089 | 2.91E-06 | -1.9547104 |
| Tbc1d2 | 1.26E-05 | -2.2142341 |
| Fcho1 | 1.85E-05 | -2.4954128 |
| Iglc2 | 3.00E-06 | -5.1136545 |
| Xist | 8.08E-17 | -5.6694142 |

**D**

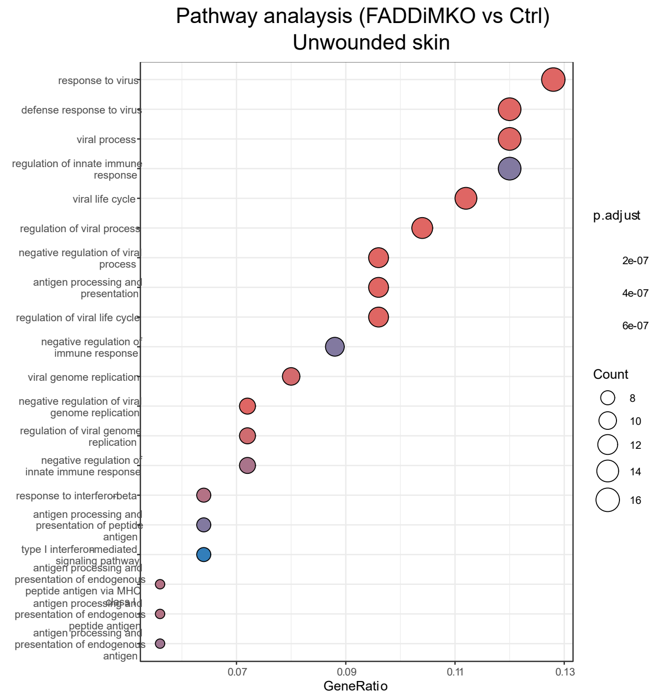


**E**

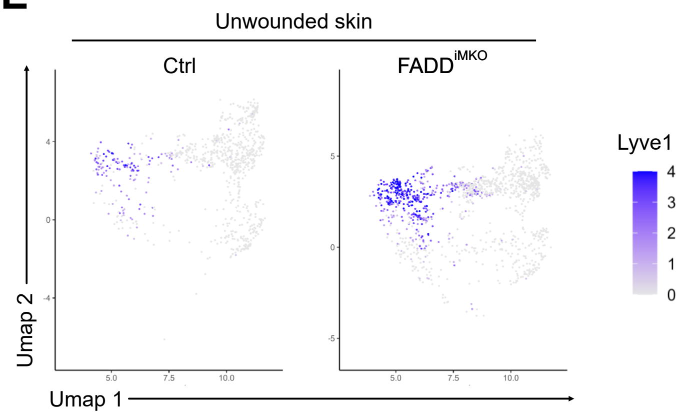
